# Emergence of Zika virus: Direct reversion of mutations and fitness restoration prior to spread to the Americas

**DOI:** 10.1101/2020.06.29.179150

**Authors:** Jianying Liu, Yang Liu, Chao Shan, Bruno T.D. Nunes, Ruimei Yun, Sherry L. Haller, Grace H. Rafael, Sasha R. Azar, Clark R. Andersen, Kenneth Plante, Nikos Vasilakis, Pei-Yong Shi, Scott C. Weaver

**Author notes:** These authors contributed equally (Jianying Liu, Yang Liu).

## Abstract

Mosquito-borne viruses have recently spread globally, with major impacts on human health. Zika virus (ZIKV) emerged from obscurity in 2013 to spread from Asia to the South Pacific and the Americas, where millions of people were infected. For the first time, severe clinical manifestations, including Guillain Barré syndrome and defects to the fetuses of pregnant women, were detected. Phylogenetic studies have shown that ZIKV evolved in Africa and later spread to Asia, and that the Asian lineage is responsible for the recent epidemics. However, the reasons for the sudden emergence of ZIKV remain incompletely understood. Accumulating evidence on other arboviruses like chikungunya and West Nile suggest the likelihood that viral mutations could be determinants of change in ZIKV transmission efficiency responsible for efficient spread. Using evolutionary analyses, we determined that four mutations, which occurred just before ZIKV introduction to the Americas, represent direct reversions of previous mutations that accompanied spread many decades ago from ZIKV’s native Africa to Asia and early circulation there. Experimental infections of mosquitoes, human cells, and mice using ZIKV strains with and without these mutations demonstrated that the original mutations reduced fitness for urban transmission, while the reversions restored fitness, increasing epidemic risk. Overall, our findings include the three newly identified, transmission-adaptive ZIKV mutations, and demonstration that these and one identified previously restored fitness for epidemic transmission soon before introduction into the America. The initial mutations may have followed founder effects and/or drift when the virus was introduced into Asia, or could be related to changes on host or vector utilization within Asia.

---

Zika virus (ZIKV), an arthropod-borne virus (arbovirus) in the genus *Flavivirus*, discovered in 1947^1^, remained obscure with little association with human disease until 2007. Then, small outbreaks occurred in Gabon^2^ and Yap Island in Micronesia^3^, associated with mild dengue-like illness. In 2013, ZIKV spread to the South Pacific, where approximately half of the residents of French Polynesia were infected^4^, and an association with Guillain Barré syndrome (GBS) was discovered^5^. Spread to other areas of the South Pacific followed; then the first outbreak ever detected in the Americas was identified in 2015 northeastern Brazil^6^, accompanied by a dramatic increase in the incidence of microcephaly^7^ and other congenital malformations now termed congenital Zika syndrome (CZS)^8^. ZIKV continued to spread to nearly all countries in the Americas with the continued association with GBS and CZS, leading the World Health Organization to declare in 2016 a Public Health Emergency of International Concern.

Early phylogenetic studies combined with ZIKV detections and seroprevalence indicated that the virus originated in and remains widespread in Africa, and was introduced many decades ago to Asia^9^. However, the reason(s) for its recent, dramatic spread to the Americas remain enigmatic. One hypothesis is that ZIKV increased its ability to be transmitted efficiently by mosquito vectors through adaptive mutations. This mechanism has been demonstrated for other arboviruses including West Nile^10^, Venezuelan equine encephalitis^11^ and chikungunya (CHIKV) viruses^12^. RNA arboviruses exhibit high mutational frequencies due to their lack of proof-reading during genome replication, providing the opportunity for rapid adaptation to changing infection and transmission opportunities^13^. The first evidence supporting this adaptive evolution hypothesis for ZIKV came from studies of an A188V amino acid substitution in the nonstructural protein 1 (NS1-A188V) that occurred just before ZIKV spread to the South Pacific and the Americas. This substitution slightly enhances ZIKV transmission by *Aedes* (*Stegomyia*) *aegypti mosquitoes*^14^, the principal epidemic vector^15^. Another substitution, V473M in the envelope protein, increases viremia in nonhuman primates, also suggesting enhanced fitness for transmission^16^. Capsid substitution T106A and several others were previously studied for their effects on mouse virulence, but none was identified. Polyprotein substitution S139N (=prM S17N) was shown to enhance murine infectivity in human and mouse neural progenitor cells and to cause more severe microcephaly in mice^17^, phenotypes not associated with more efficient transmission via viremia.

## Directly reverting Zika virus mutations

To further test the hypothesis that adaptive evolution enhanced epidemic ZIKV transmission, leading to unprecedented spread, millions of infections, and the resulting detection of rare disease manifestations, we performed phylogenetic analyses to identify additional amino acid substitutions that preceded its spread from Asia to the Americas. Extended Data Fig. 1 shows a tree with four substitutions mapped to branches that preceded epidemic spread, including NS1-A188V^14^. Tracings of these amino acids in ZIKV phylogenetic trees are shown in Extended Data Fig. 2. Strikingly, when we traced the evolution of these 4 pre-epidemic amino acid substitutions, we found that they all represented direct reversions of earlier mutations inferred to have accompanied spread from Africa to Asia, or early circulation in Asia. None of these initial mutations was noted in earlier, comprehensive phylogenetic studies^18^. The simplest explanation for this mutation pattern is that the pre-epidemic substitutions restored fitness (reproduction and survival ability) declines caused by founder effects and/or genetic drift during early, inefficient *A. aegypti-borne* interhuman transmission. This transmission inefficiency is consistent with the lack of recognized outbreaks in Asia before 2010, and bottlenecks that accompany the arbovirus transmission cycle can allow drift to dominate evolution if populations remain small.

Founder effects result in a loss in genetic diversity that accompanies geographic introductions of small founder populations from a large, ancestral population. For human-amplified arboviruses like ZIKV, these typically involve a single infected traveler^19^. Furthermore, several virus population bottlenecks are known to punctuate the arbovirus transmission cycle during mosquito infection and dissemination of virus to the salivary glands. The resultant loss of genetic diversity can result in the fixation of random mutations, the majority of which are, by chance, deleterious for RNA viruses^20^ and other organisms. The hypothesis that African ZIKV strains lost fitness upon their introduction into Asia is also supported by their greater infectivity for *A. aegypti* mosquitoes compared to Asian and subsequent American ZIKV strains^21–23^, as well as their greater virulence in mouse models of human infection^23,24^.

As an initial test of the hypothesis that ZIKV underwent a fitness decline upon its introduction into Africa, followed by fitness restoration, we assessed the fitness of African, Asian, and American ZIKV strains. We used competition fitness assays, where ZIKV strains differing by as little as one amino acid were generated from cDNA clones and mixed in a roughly 1:1 ratio based on Vero cell plaque-forming units (PFU). This inoculum mixture was then used to initiate infections. Following appropriate incubation to allow the two strains to compete for replication and, in mosquitoes, dissemination to the salivary glands, the virus mixture was again quantified by Sanger sequencing of RT-PCR amplicons to estimate the ratios of mixed nucleotides using electropherogram peaks. The relative fitness value w was analyzed according to w=(f0/i0), where f0 is the final ratio of one competitor following infection, and i0 is the initial ratio in the inoculum or bloodmeal mixture; this ratio reflects relative fitness advantage of one competitor over the other (Fig. 1 a). This approach has major advantages over performing individual strain infections with numerous host replicates; each competition is internally controlled, eliminating host-to-host variation that can reduce the power of experiments, and the virus strain ratios can be assayed with more precision than individual virus titers. Thus, competition assays have been used for many studies of microbial fitness^25^, including arboviruses^26–28^. Because the competition assay relies on electropherogram peak measurements from Sanger sequencing to quantify mutant ratios, with the potential for inconsistency, we conducted experiments to validate the consistency and accuracy of this method. Extended Data Fig. 3 shows the very high degree of accuracy and consistency of this method. To confirm that our targeted 1:1 ratios were similar in mosquito experimental systems, where RNA:infectious virus ratios could differ based on different host-dependent levels of mutant infectivity, compared to those for Vero cells, we also determined infectious titers of the wt and mutants on C6/36 mosquito cells, and compared them with the Vero PFU ratios. These ratios were nearly indistinguishable with no significant differences (Extended data Fig. 4), indicating that frequency-based differences in fitness based on potential mutation effects on initial infectivity were not a concern.

**Figure 1.**
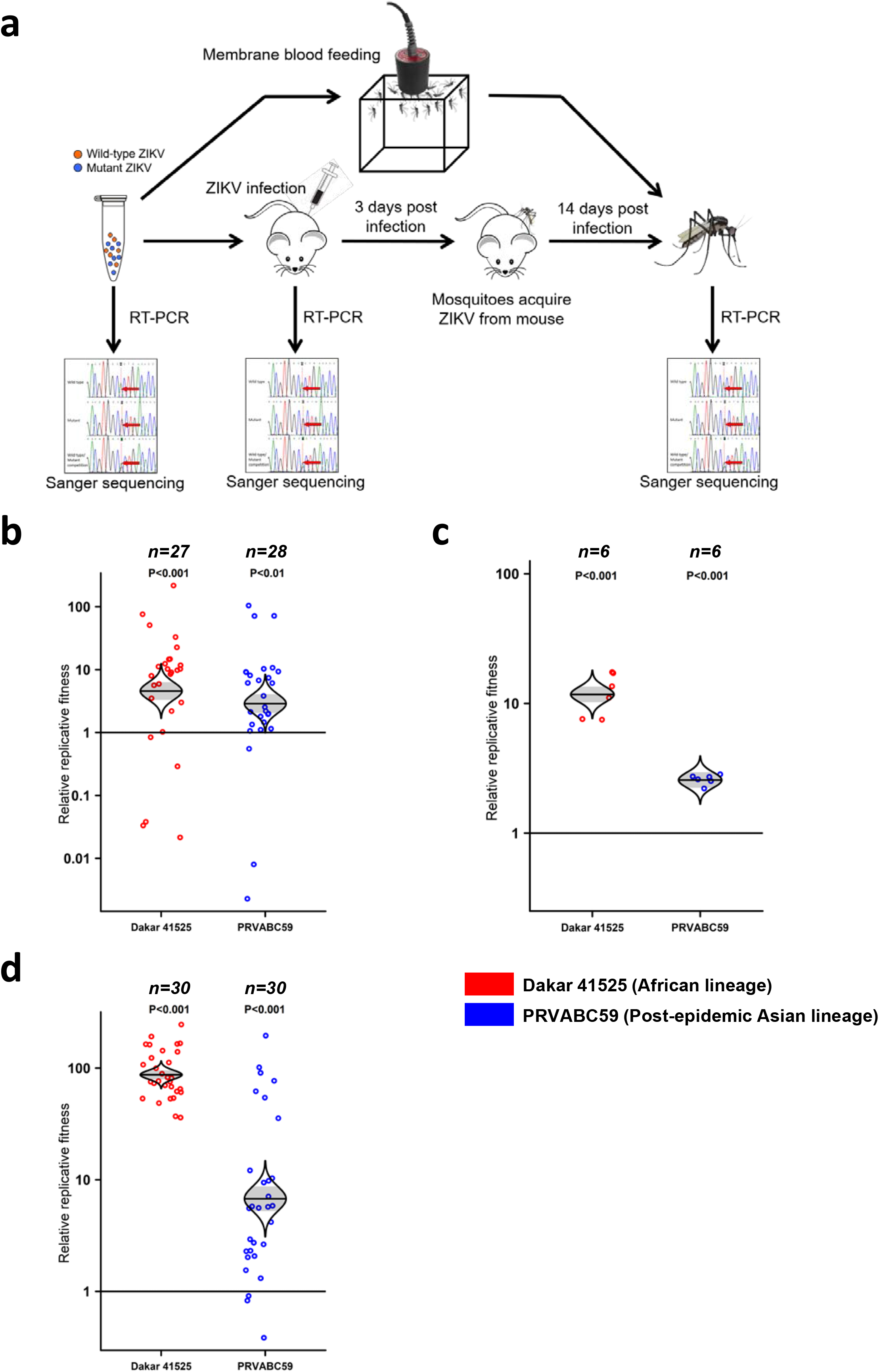
African lineage ZIKV and post-epidemic Asian lineage ZIKV had the fitness advantage versus pre-epidemic Asian lineage ZIKVs in both A129 mice and mosquitoes. **a**, Experimental design of competition fitness assays. African lineage ZIKV and post-epidemic Asian lineage ZIKV were mixed with pre-epidemic Asian lineage ZIKV at an approximate ratio of 1:1 respectively. The initial 1:1 ratios of mixed virus were confirmed by RT-PCR, followed by Sanger sequencing of replicons and polymorphic nucleotide peaks analysis. The ZIKV mixture was engorged by mosquitos through membrane blood feeding or biting an infected A129 mouse at the viremia peak 3 days post-infection. The mosquitoes were sacrificed after 14 days of incubation, the ZIKV population amplified by RT-PCR, and the amplicon Sanger-sequenced. The blood and organs of infected mice were collected 3- or 8-days post-infection, respectively. All the mosquitoes and mouse specimens were subjected to RT-PCR amplification and Sanger sequencing to compare the ZIKV strain ratio after competition, and each point represents a single mosquito or mouse sample. Relative fitness values were compared among multiple competition replicates to determine if fitness of the competitors differed. **b**, Fitness comparison between African (Dakar 41525), Asian pre-epidemic (FSS13025) and American (PRVABC59) ZIKV strains in *A. aegypti.* The relative replicative fitness value w was analyzed according to w=(f0/i0), where i0 is the initial ratio of the Dakar 41525 or PRVABC59 competitor to the FSS13025 strain, determined from Sanger sequence electropherogram peaks, and f0 is the final ratio. **c**, Fitness comparison of African (Dakar 41525), Asian pre-epidemic (FSS13025) and American (PRVABC59) ZIKV strains in mouse blood, using the same method as for panel b. **d**, Fitness comparison of African (Dakar 41525), Asian pre-epidemic (FSS13025) and American (PRVABC59) ZIKV strains in mosquitoes after feeding on the viremic mice shown in panel c, and extrinsic incubation, using the same method as for panel b. **b**-**d**, The distribution of the model-adjusted means is illustrated by catseye plots with shaded +/− standard errors overlaid by scatterplots of individual relative fitness values; scatterplots have been randomly jittered horizontally for clarity, and are shown on the log (base-10) scale such that comparisons are against a null value of 1.

## Fitness of major Zika virus lineages

Initially, we performed competition assays with wildtype (*wt*) ZIKV strains representing the African, Asian pre-epidemic, and American epidemic genotypes (Fig. 1). The highly susceptible Rockefeller strain of *A. aegypti* mosquitoes was used along with interferon type I receptor-deficient A129 mice, models for human infection^29,30^. Following infection with an approximately 1:1 mixture of the African Dakar 41525 and Asian Cambodian FSS13025 strains, or an equivalent mixture of FSS130025 and the Puerto Rican RRVABC59 strain, samples from mosquito bodies and mice were evaluated for changes in the initial ratio, indications of competitive fitness (Fig. 1a). Following oral mosquito infection and incubation, the African strain consistently and significantly won the competition with the Asian strain, and the Puerto Rican strain consistently outcompeted the Asian strain (Fig. 1b). These findings were reproducible when additional African, Asian and American ZIKV strains, as well as a Dominican Republic strain of *A. aegypti* with a low level of colonization (F6), were used (Extended Data Fig. 5). After three days of competition following subcutaneous infection of A129 mice to mimic human infection, the same relative fitness results were obtained when viremic blood (the vertebrate host compartment where major selection for efficient mosquito infection occurs) was analyzed (Fig. 1c). Analysis of individual organ samples from mice on day 8 (after viremia had subsided to eliminate this confounding source of virus in organs) showed that these competition outcomes were consistent for replication throughout the mouse (Extended Data Fig. 6). Mosquitoes that fed upon these viremic mice were also evaluated after extrinsic incubation, with the same results of greater fitness of the African versus Asian ZIKV strain, and greater fitness of the American versus Asian strain (Fig. 1d).

Next, primary human cells believed to be involved in seeding viremia were used for competition assays: fibroblasts and keratinocytes^31,32^. The mixed ZIKVs were inoculated into cells and the culture supernatant were harvested, RT-PCR-amplified and Sanger sequenced 3 days post-infection (Extended Data Fig. 7a). The results showed that the African strain and post-epidemic American strains always out-competed the pre-epidemic Asian strain in both cell types (Extended Data Fig. 7b, c). In all of these experimental models, the changes in competitor ratios appeared to be greater between the African versus Asian strains compared to the Asian versus American strains. Overall, these data supported the hypothesis that ZIKV underwent a major loss of fitness upon its introduction into Asia, followed by a partial restoration of fitness upon introduction into the Americas. We note that microcephaly and other central nervous system involvement is rare and not known to play a major role in generating viremia, so there should not be major selection for CNS tropism to enhance viremia and subsequent transmission by mosquitoes. Therefore, microcephaly may represent a chance pathogenic outcome that did not result from positive selection. Also, sexual ZIKV transmission is believed to play a minor role compared to vector-borne transmission.

Based on the evidence discussed above, we developed a revised adaptation hypothesis, depicted in Fig. 2a, that the 4 initial amino acid substitutions (and possibly others not undergoing reversion) represent founder effects that reduced the fitness of ZIKV for transmission in the epidemic cycle by reducing *A. aegypti* transmissibility and/or human viremia levels. The reversion of these 4 mutations, and possibly other adaptive mutations, then partially restored ZIKV fitness, allowing for efficient spread to the Americas and major epidemics.

**Figure 2.**
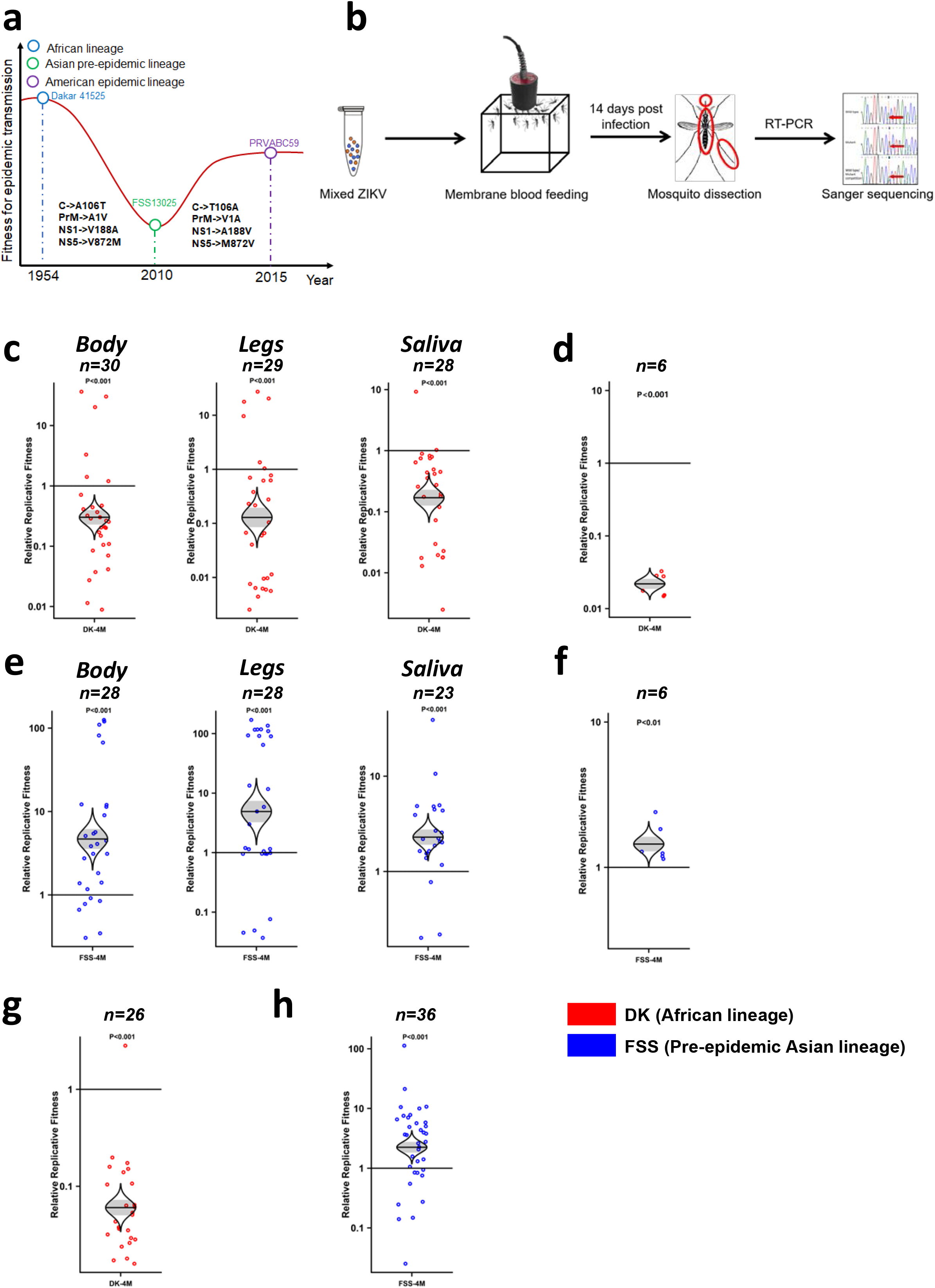
Four amino acids determine much of the fitness difference between different ZIKV strains in both mosquitoes and A129 mice. **a**, The hypothetical diagram of fitness changes following the spread of ZIKV from Africa to Asia and the Americas, including strains used for fitness assays. **b**, Schematic representation of competition fitness assay in mosquito bodies, legs and saliva. **c**, **e**, Fitness comparison between African and Asian lineage, and the four amino-acid substitution mutants (DK-4M and FSS-4M) in mosquito bodies, legs and saliva. The mosquitoes acquired the ZIKV mixture through membrane blood feeding and were assayed at 14 days post-infection. The body (carcass), legs and saliva were collected from each individual mosquito and subjected to RT-PCR amplification and Sanger sequencing to determine relative fitness values of 4M over wt; each point represents a single mosquito or mouse sample. **d**, **f**, Competition between DK-WT and DK-4M(**d**), FSS-WT and FSS-4M (**f**) in mouse blood collected 3 days after infection. **g**, **h**, Fitness of DK-4M (**g**) and FSS-4M (**h**) in mosquitoes after oral infection from viremic mice. The mosquitoes acquired the virus through biting a viremic mouse 3 days after infection and were tested after 14 days of extrinsic incubation. **c**-**h**, The distribution of the model-adjusted means is illustrated by catseye plots with shaded +/− standard error bars overlaid by scatterplots of subject measures; scatterplots have been randomly jittered horizontally for clarity, and are shown on the log (base-10) scale such that comparisons are against a null value of 1.

## Fitness effect of reversion mutations

To test this hypothesis that the 4 revertant mutations restored ZIKV fitness, we used *A. aegypti* mosquitoes and A129 mice in more detailed studies of the combinations of four amino acid substitutions that underwent reversion. For mosquito infections, we assayed bodies to evaluate infection and initial replication, legs to sample the hemolymph, which contains virus that has disseminated from the digestive tract (midgut) into the hemocoel for access to the salivary glands, and saliva collected in vitro to assess the virus population available for transmission (Fig. 2b). Competition between the African ZIKV strain and a mutant containing the four initial amino acid substitutions resulted in a consistent advantage for the African strain in all mosquito samples (Fig. 2c), as did infection of A129 mice as sampled in major organs and blood (Fig. 2d, Extended Data Fig. 8a). In contrast, when the four reversion substitutions were placed into the pre-epidemic Asian strain to recapitulate ZIKV evolution just before introduction into the Americas, these substitutions resulted in a major fitness gain (Fig. 2e, f, Extended Data Fig. 8b). When mosquitoes were fed upon the viremic mice, the same outcome was observed (Fig. 2g, h). These results indicate that the four substitutions that directly reverted prior to the introduction of ZIKV into the Americas were major components of the fitness differences among African, Asian and American virus strains. At the same time, similar competition assays were performed in human fibroblasts and keratinocytes. The ZIKV strains containing these four initially substituted amino acids always won the competition in these human primary cells (Extended Data Fig. 9).

To assess the roles of the individual amino acid substitutions in the overall fitness differences among ZIKV strains, we tested them individually using the same experimental systems. Placed individually into the African strain, all four initial mutations reduced overall fitness for infection, dissemination (two mutants showed no fitness effect in this compartment) and transmission by mosquitoes, although only C-A106T and NS1-V188A results were significant (Fig. 3a-c). The same trends were observed in mosquito bodies, legs and saliva (Fig. 3a-c). When the reversions were placed into the Asian strain, only the NS1-A188V mutant produced a significant increase in ZIKV infection in the mosquito bodies, legs and saliva, indicating this amino acid may have a dominant role during the competition (Fig. 3d-f). As with the 4X mutants, the lack of detection of one competitor in saliva samples indicates a major fitness advantage for vector transmission. To confirm fitness effects in models for the vertebrate amplification host, the Dakar NS1-V188A and FSS NS1-A188V mutants were mixed with corresponding *wt* strains and inoculated into A129 mice. The ZIKVs containing NS1-226V exhibited a fitness advantage in both mice (Fig. 3g, h, Extended Data Fig. 10) and mosquitoes (Fig. 3i, j). The slight asymmetry in the fitness effects between some of the initial 4 mutations in the African strain and the reversions in the Cambodian pre-epidemic strain could reflect epistatic interactions that differ between the two virus strains.

**Figure 3.**
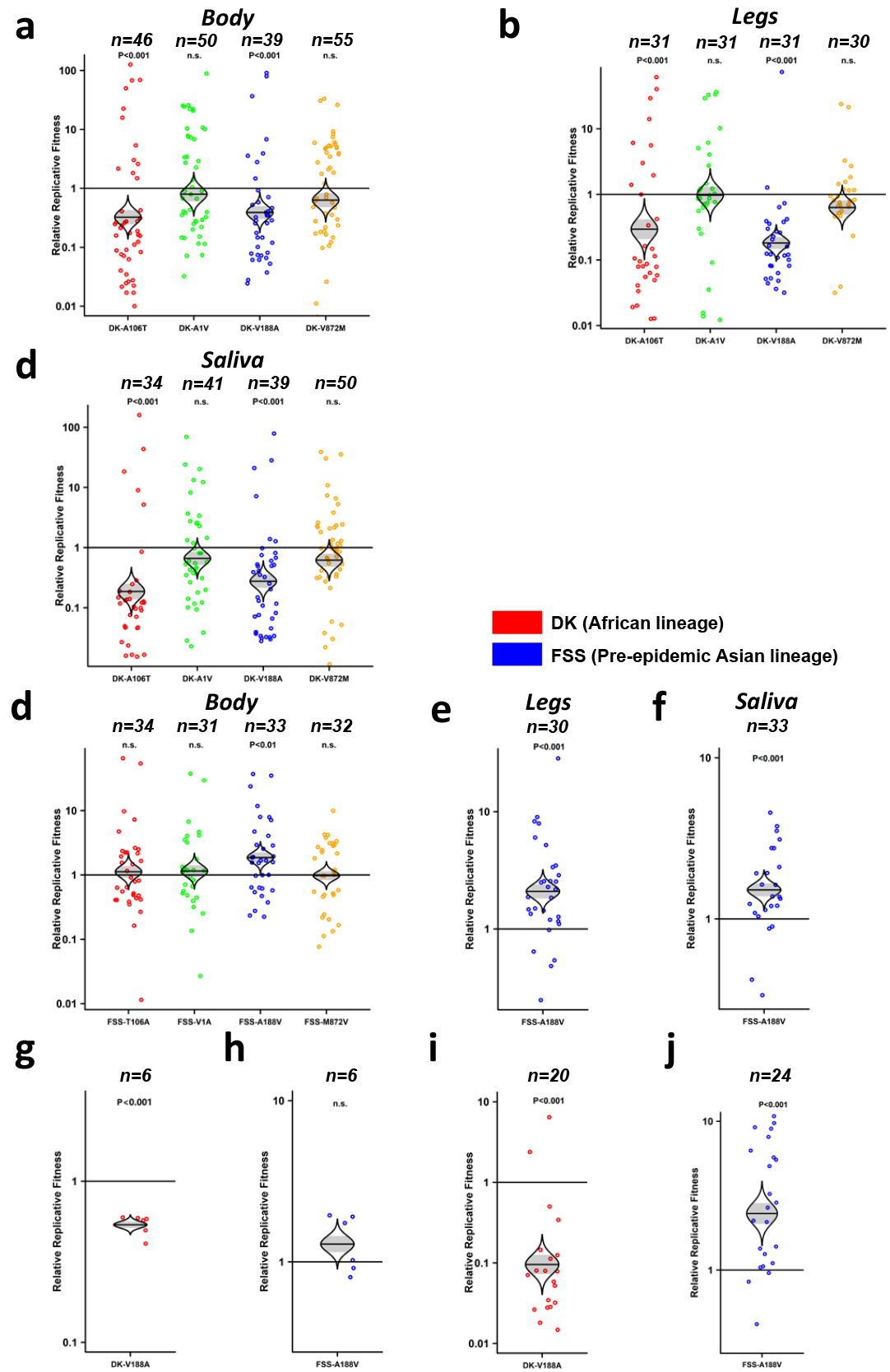
The fitness comparison of 4 individual mutant viruses against wild-type ZIKV strains in mosquitoes and A129 mice. **a**-**c**, The relative fitness of individual mutants in the Dakar strain, tested in mosquito bodies (**a**), legs (**b**) and saliva (**c**). **d**-**f**, The fitness comparison of individual reversion mutants in the FSS13025 strain, tested in mosquito bodies (**d**), legs (**e**) and saliva (**f**). **g**,**h**. Each point represents a single mosquito sample. The fitness comparison of DK-V188A (**g**) and FSS-A188V (**h**) versus wild-type strains in mouse blood. **i**, **j**, The fitness comparison of DK-V188A (**i**) and FSS-A188V (**j**) in mosquitoes after oral infection from viremic mice. **a**-**j**, The distribution of the model-adjusted means is illustrated by catseye plots with shaded +/− standard error overlaid by scatterplots of subject measures; scatterplots have been randomly jittered horizontally for clarity, and are shown on the log (base-10) scale such that comparisons are against a null value of 1.

Finally, the overall fitness of the sets of four mutations or reversions were tested over a complete experimental transmission cycle. To initiate the cycle, mosquitoes were inoculated intrathoracically (to ensure uniform infection) with mixtures of the African ZIKV strain and the four initial mutations, or the Asian strain with the four reversions (Fig. 4a). Following six days of extrinsic incubation (mosquitoes become infectious more rapidly after intrathoracic than oral infection), these mosquitoes were exposed to A129 mice for oral transmission. Three days later during peak viremia, naïve mosquitoes were allowed to feed on the mice, then these mosquitoes were assayed after 14 days of extrinsic incubation as in the above experiments. In salivary glands following incubation after mosquito inoculation, the initial four mutations decreased the fitness of the African ZIKV strain (Fig. 4b), and the four reversions increased the fitness of the Asian strain (Fig. 4c) as observed after oral infection (Fig. 2c, e). These fitness effects continued during infection of mouse blood and organs (Fig. 4d, e, Extended Data Fig. 11) and also in mosquitoes fed orally on these mice during viremia (Fig 4f, g). Overall, these results demonstrate that the four initial mutations reduced the fitness of the African ZIKV strain through a complete transmission cycle, while the four reversions at least partially restored fitness.

**Figure 4.**
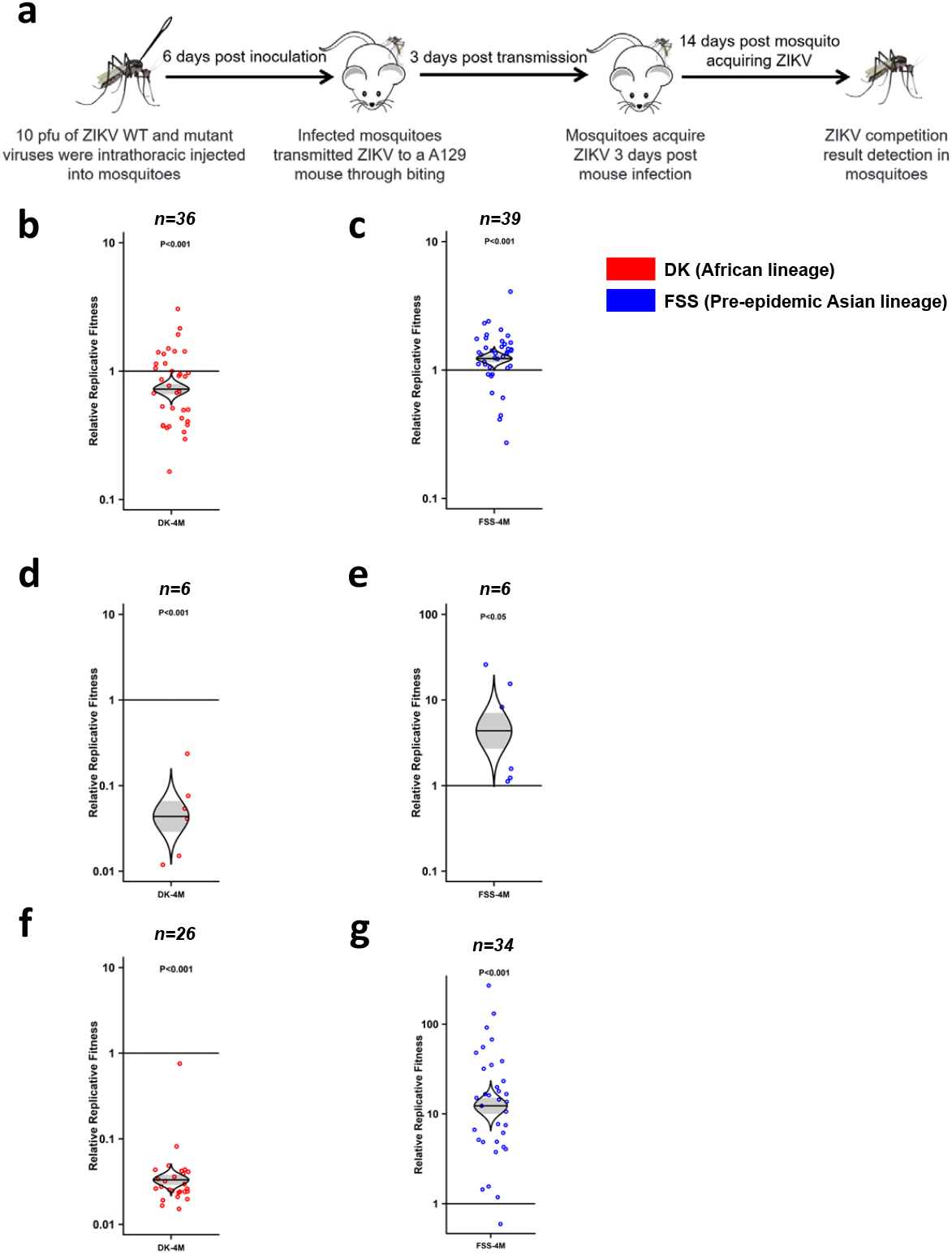
Fitness comparison of DK-4M and FSS-4M against wild-type ZIKV strains during a mosquito–host–mosquito transmission cycle. **a**, Schematic representation of the study design. 10 pfu of ZIKV WT and mutant viruses were intrathoracically injected into mosquitoes. After 6 days of extrinsic incubation, the infected mosquitoes fed on a A129 mice to transmit virus. Naïve mosquitoes were then employed to bite the infected A129 mice 3 days post-mouse infection. After an additional 14 days, the mosquitoes were harvested. All the mosquitoes and mice samples collected during the transmission cycle were analyzed by RT-PCR and Sanger sequencing. Each point represents a single mosquito or mouse sample. **b**, **c**, Fitness comparison of DK-4M (**b**) and FSS-4M (**c**) against wild-type viruses in mosquito salivary glands 6 days post-intrathoracic injection. **d**, **e**, Fitness of DK-4M (**d**) and FSS-4M (**e**) versus wild-type viruses in mouse blood. **f**, **g**, The fitness comparison of DK-4M (**f**) and FSS-4M(**g**) versus wild-type viruses in mosquitoes after oral infection from viremic mice. **b**-**g**, The distribution of the model-adjusted means is illustrated by catseye plots with shaded +/− standard error overlaid by scatterplots of subject measures; scatterplots have been randomly jittered horizontally for clarity, and are shown on the log (base-10) scale such that comparisons are against a null value of 1.

## Evolution of ZIKV fitness in Asia

In summary, we identified three ZIKV amino acids (C-106, prM-1 and NS5-872) that we show for the first time affect virus transmission by increasing infection and transmission efficiency by *A. aegypti* and/or enhancing replication in models for human infection. We also demonstrate for the first time that, like those three residues, the NS1-A188V, shown earlier to enhance vector infection^14^, also represents a direct reversion of a mutation that occurred soon after ZIKV reached Asia. Although all South Pacific and American ZIKV strains include all four reversions, some recent Asian strains also have some of these revertant residues. This suggests that the reversions did not all occur immediately before ZIKV introduction to these epidemic locations, which is consistent with emergence involving a combination of adaptive mutations that predispose virus strains to epidemic initiation, along with stochastic factors related to chance introductions into naïve, epidemic-permissive locations. Also, the lower fitness of the 4X reversion mutant compared to the American strains (Fig. 2) indicates that other mutations that do not represent reversions were likely also involved in emergence. These conserved amino acid substitutions in distal tree branches and their possible phenotypes are listed in Extended Data Table 1.

Overall, our results support the hypothesis that ZIKV underwent fitness declines upon its introduction into Asia many decades ago, and/or during its initial stages of circulation there when a lack of recognized outbreaks suggests limited urban circulation. An alternative hypothesis that would not rely on drift is that ZIKV host range changed upon its introduction into Asia, followed by another change (or reversion in host usage) just before introduction into the Americas; both of these hypothetical host range changes could have resulted in positive selection that involved the same four amino acids, resulting in reversion. The simplest form of this hypothesis would be that a sylvatic cycle involving different vectors and vertebrate hosts from those in Africa was the initial form of Asian transmission, and the urban cycle only developed there relatively recently accompanied by the four reversions. Although there is no evidence of enzootic ZIKV transmission in Asia, and the 1966 ZIKV isolation from *A. aegypti* mosquitoes in Malaysia suggests that human-amplified transmission has been ongoing in Asia for many decades, the detection of a sylvatic cycle is difficult in the absence of extensive genetic divergence from urban strains.

## Parallels with CHIKV

The hypothesis that the initial urban ZIKV cycle was inefficient due to founder effects and drift, is consistent with the same pattern of fitness loss that accompanied CHIKV introduction into Asia about a century ago, when a major deletion in the 3’ untranslated genome region occurred. This fitness of this Asian lineage remains incompletely restored to this day^31^. However, the inability of the four ZIKV reversion mutations to generate fitness comparable to that of African strains suggests that additional mutations, possibly also the result of founder effects or drift but without direct reversions, also limited transmission and spread in Asia. Additional mutations that occurred during circulation in Asia must be tested to expand our results to a comprehensive understanding of ZIKV evolution, and understanding the mechanisms of fitness effects of the four substitutions we studied require further study.

These results as well as the evolutionary history of ZIKV have remarkable parallels to those of CHIKV, which also evolved in Africa and spread many decades ago to Asia before its recent introduction into the Americas^12^. Previous studies indicate that the CHIKV arrived in Asia many decades ago with debilitating mutations in its 3’ untranslated genome region (3’UTR), followed by only partially fitness restoration over many decades through point mutations and a duplication^33^. Another mutation in the E1 glycoprotein, also a probable founder effect (no phenotype has been established for this mutation on its own), prevented for many decades CHIKV adaptation for efficient transmission by *A. albopictus* mosquitoes in Asia, their native territory^34^. Then, just before or upon its introduction into the Americas in 2013, a duplication in the 3’UTR also improved CHIKV fitness^35^. However, unlike ZIKV, another lineage of CHIKV has also undergone a series of adaptive mutations to enhance its ability to use *A. albopictus* for transmission, with no evidence of founder effects, and the relative fitness advantages of these CHIKV mutations far exceed those of the four ZIKV mutations that we studied; for example, individual *A. albopictus-adaptive* CHIKV mutations showed relative fitness values of 5-40, while even the combination of 4 ZIKV mutations we studies had a lower overall fitness effect than any of these individual CHIKV mutations (Extended Data Table 2). The similarities in the presumed roles of founder effects in the evolution and emergence potential of ZIKV and CHIKV suggest that drift has been understudied as a factor in the emergence of arboviral and other RNA viral diseases, and that the stochastic nature of founder effects may limit our ability to ultimately predict the emergence of new viral diseases.

Finally, the high fitness of the African ZIKV strains in all experimental systems we utilized raises important questions about the risk of outbreaks and severe disease on that continent. The greater transmissibility of these African strains, even compared to American strains^21^, raises the question of why outbreaks have never been detected in Africa aside from one in Angola caused by a strain imported from the Americas^36^. Limited African surveillance and the widespread presence of other infections that typically lead to Zika misdiagnosis in the absence of laboratory diagnostics could be responsible. Herd immunity in Africa, which could also play a role in limiting outbreaks as well as the incidence of congenital Zika syndrome, has been reported to range from 1-52%^37^. However, some studies have used methods other than neutralization that could be highly cross-reactive with other endemic flaviviruses. We also cannot rule out the possibility that our models for human infection are not suitable to determine ZIKV fitness for human amplification. Additional surveillance and work with more human cells and possibly nonhuman primate models are needed to further explore this possibility and to better understand the lack of Zika outbreaks in Africa caused by African lineage strains.

In addition to the fitness for transmission that we examined, the history of ZIKV’s ability to cause GBS and CZS requires further study. It is possible that enhanced viremia just before spread to the Americas, suggested by our model systems involving human cells and A129 mice, could at least partially explain the emergence of these severe disease outcomes if viremia magnitude is correlated with infection of the placenta. However, better surveillance in Africa is ultimately needed to determine if American or recent Asian strains are unique in this pathogenic potential.

## Methods

Mouse and mosquito studies were performed in accordance with the guidance for the Care and Use of Laboratory Animals of the University of Texas Medical Branch (UTMB). The protocol (protocol number 1708051 for mouse) were approved by the Institutional Animal Care and Use Committee (IACUC) at UTMB. All the mouse manipulations were performed under anesthesia by isoflurane.

### Mice, mosquitoes, cells and viruses

A129 mice, which are deficient in type I interferon receptors, were bred and maintained in animal biosafety level 2 (ABSL2) facilities at UTMB. Sex-matched mice, randomly selected and 6-to-8-week-old, were used. The *Aedes aegypti* Rockefeller strain and *A. aegypti* Dominican Republic strain (F6) were maintained in an incubator at 28°C and 80% humidity. Vero cells (CCL81) and C6/36 cells for virus rescue, proliferation and titration were purchased from the American Type Culture Collection (Bethesda, MD, USA) and maintained in Dulbecco’s modified Eagle’s medium (Gibco, Waltham, MA, USA) with 10% heat-inactivated fetal bovine serum (Atlanta Biologicals, Flowery Branch, GA, USA), and supplemented with 1% Tryptose phosphate broth (Gibco, Waltham, MA, USA) for C6/36 cells. The human primary dermal fibroblast cells and human epidermal keratinocyte cells were purchased from Lonza (Walkersville, MD, USA) and were maintained in FGM-2 BulletKit (Lonza) and KGM-Gold BulletKit (Lonza) media, respectively. All cells were verified and tested negative for mycoplasma. The parental African lineage ZIKV Dakar 41662 strain (KU955592), and 41671 stain (KU955595), and Asian lineage ZIKV Dominican Republic R114916 (KX766028), and Honduras HN-ME 59 stains were obtained from the World Reference Center for Emerging Viruses and Arboviruses at UTMB. The ZIKV DK-WT (Dakar wild-type 41525 strain, KU955591), DK-A106T (capsid 106th amino-acid), DK-A1V (pre-membrane 1st amino-acid), DK-V188A (nonstructural protein 1 188th amino-acid), DK-V872M (nonstructural protein 5 872nd amino-acid) were constructed using standard site-directed mutagenesis of cDNA clones and rescued on Vero cells using electroporation of transcribed RNA. The FSS-WT (FSS13025 wild-type strain, KU955593), FSS-T106A, FSS-V1A, FSS-A188V and FSS-M872V, as well as reversion mutants placed in the ZIKV Dakar strain, were constructed in the same manner, along with the DK-4M and FSS-4M that combined all four mutations on the matched backbone.

### Competition assay

To better understand the relative fitness of different ZIKV strains and mutants, we conducted competition assays between two virus strains or a wild-type and mutant strain in various models. Initial 1:1 virus:mutant mixtures were made based on PFU titers determined in Vero cells, and virus ratios used to calculate fitness were determined by Sanger sequencing of RT-PCR amplicons. The PFU:genome ratio was consistent among all of our ZIKV strains and constructs, as estimated from the inocula and bloodmeals by comparing Vero cell PFU to genome copies as estimated from real-time RT-qPCR standard curves. To confirm that ratios were also similar in mosquito experimental systems, where RNA:infectious virus ratios could differ based on different levels of infectivity compared to Vero cells, we also determined infectious titers of the wt and mutants on C6/36 mosquito cells, and compared them with the Vero-PFU ratios.

### Mosquitoes

The *Aedes aegypti* Rockefeller strain and *A. aegypti* Dominican Republic strain (F6) were used for the following studies. Triplicate mixtures of viruses were diluted to 6 log_10_ PFU/ml mixed at a 1:1 ratio based on Vero PFU titers. Virus mixtures were subsequently mixed at a ratio of 1:1 with PBS-washed sheep blood cells (Hemostat laboratories, Galveston, TX, USA). Aliquots of blood meals were collected to verify the initial virus proportions in the inoculum. Following a 14-day extrinsic incubation, saliva expectorated into capillary tubes, legs representing virus disseminated into the hemocoel, or remaining mosquito bodies were collected separately in a 2ml Eppendorf Safe Lock tube with 250μl DMEM (2% FBS, 1% Sodium Pyruvate) with 25 mg/ml Amphotericin B and a stainless steel bead (Qiagen, Hilden, Germany). Samples were stored at −80°C.

### A129 mice

Sex-matched A129 mice (6-8 weeks old) were used for ZIKV competition assays. The A129 mouse is defective for the interferon type I receptor and susceptible to ZIKV infection in vivo with high titered viremia. Mixtures of ZIKV strains (total 1×10^4^ PFU/mouse) were inoculated intradermally to simulate a mosquito bite. Titers of viruses before mixing were plaque assayed to verify proper concentrations. Also, an aliquot of the inoculum was reserved for estimating the initial ratio of viruses. Mice were bled retro-orbitally on day 3 post infection representing the peak. Serum was collected from blood via centrifugation at 2,000 g for 5 minutes and stored at −80°C until analysis. Eight days post infection, all infected mice were euthanized and necropsied and major organs collected in a 2ml Eppendorf Safe Lock tube with 500μl DMEM (2% FBS, 1% Sodium Pyruvate) and a stainless-steel bead. Samples were stored at-80°C.

### Human primary cells

The human fibroblasts and keratinocytes were purchased and qualified by ATCC. Cells were seeded into 12-well plates 1 day before infection to allow them reach 90% confluence. Mixtures of ZIKVs (1 PFU/cell MOI) were added into the cells and incubated at 37°C for 2 hours. Titers of viruses before mixing were plaque assayed to verify proper concentrations. Also, an aliquot of the inoculum was reserved for estimating the initial ratio of the viruses. After incubation, the cells were washed 3 times with 1ml of PBS. The infected cells were cultured, and the supernatant were collected from days 1-5. Samples were stored at −80°C for further analysis.

### Sample preparation and Nucleic acid extraction

Aliquots of mosquito lysates, mouse serum, mouse organ lysates and human primary cell supernatant samples were prepared in RNeasy Mini Kit (Qiagen) lysis buffer RLT (400μl), and nucleic acids were subsequently extracted according to the manufacturer’s protocol. After extraction, a portion of the RNA was immediately applied to a one step RT-PCR assay and the remaining material was archived at −80°C.

### Quantitative real-time RT-PCR assays

Prior to Sanger sequencing analysis, virus-positive samples were identified by quantitative real-time RT-PCR, which was performed using a QuantiTect Probe RT-PCR Kit (Qiagen) on the LightCycler 480 system (Roche, Rotkreuz, Switzerland) following the manufacturer’s protocol. The primers and probes are listed in Extended Data Table 3. The absolute quantification of ZIKV RNA was determined using a standard curve with in vitro-transcribed, full-length ZIKV RNAs.

### Reverse transcriptase PCR

400-500 bp of RT-PCR product were synthesized and amplified from extracted RNA using a SuperScript™ III One-Step RT-PCR kit (Invitrogen, Carlsbad, CA, USA). 20μl reactions were assembled in PCR 8-tube strips through the addition of 10μl 2X reaction mix, 0.4μl SuperScript™ III RT/Platinum™ Taq Mix, 0.8μl 10 μM specific forward primer, 0.8μl 10 μM specific reverse primer (see Extended Data Table 2), 4μl of extracted RNA and 6μl Rnase-free water. RT was completed using the following protocol: 1) 55°C, 30 min; 94°C, 2 min; 2) 94°C, 15s; 60°C, 30s; 68°C, 1min; 40 cycles; 3) 68°C, 5 min; 4) indefinite hold at 4°C. 2μl of generated PCR products were loaded to 1% DNA agarose gel to verify the size. The left samples were purified by a QIAquick PCR Purification kit (Qiagen) according to the manufacturer’s protocol. The concentration of RT-PCR products >10 ng/ul were qualified following Sanger sequencing.

### Sanger sequencing and electropherogram peak height analysis

Sequences of the purified RT-PCR products (concentration >10 ng/ul) were generated using a BigDye Terminator v3.1 cycle sequencing kit (Applied Biosystems, Austin, TX, USA). The products of the sequencing reactions were then purified using a 96-well plate format (EdgeBio, San Jose, CA, USA), and then analyzed on a 3500 Genetic Analyzer (Applied Biosystems). The peak electropherogram height representing each mutation site and proportion of each competitor was analyzed using the QSVanalyser program^39^.

### Intrathoracic inoculation of mosquitoes

The intrathoracic microinjection of ZIKV into mosquitoes has been described previously^40^. Female mosquitoes were anaesthetized on a cold tray (0°C). Then, a defined titer (1 pfu in 100 nl total inoculum volume) of mixed ZIKV strains were inoculation into the mosquito thorax with a Nanoject III (Drummond, Pennsylvania, USA). Six days later, the infected mosquitoes were allowed to feed on A129 mice to transmit ZIKV.

### Membrane blood feeding

Seven-day-old female mosquitoes were placed into mesh-covered cartons and provided cotton balls containing 10% sucrose. Complement-inactivated sheep blood was mixed with different ZIKV combinations for mosquito feeding via a Hemotek system membrane feeder (5W1, Hemotek limited, Lancashire, UK). The total viral titers of the two competing ZIKVs in the blood meals were 1×10^6^ PFU/ml. Fully engorged mosquitoes were incubated for 14 days until harvest. At that time, live mosquitoes were killed by freezing and homogenized individually (TissueLyser II, Qiagen) for RNA isolation and real-time qPCR detection. The infected mosquito RNAs were RT-PCR amplified, followed by Sanger sequencing.

### Mosquito feeding on infected mice

Ten female mosquitoes were placed into mesh-covered cartons for blood feeding following sugar-starvation for 24 hrs. ZIKV-infected A129 mice were anaesthetized with ketamine and placed on the top of the cartons for 20 min of feeding in the dark. Then, fully engorged mosquitoes were transferred to new containers and incubated for 14 days prior to RT-PCR, followed by Sanger sequencing of amplicons.

### Plaque assay

The plaque assay was performed on Vero cells following procedures described previously^41^. In brief, Vero cells were seeded into 12-well plates 12 to 16 hours prior to the plaque assay. 25 μl of each sample were used for 10-fold serial dilutions. For each dilution, 200 μl were added to 12-well plates with 90% confluent Vero cells. The cells were incubated at 37 °C with 5% CO_2_ for 1h with gentle rocking every 15 min. After that, 1 ml of overlay (DMEM, 2% FBS, containing 0.8% methyl cellulose with 1% antibiotics) was added onto each well. The plates were cultured at 37°C with 5% CO_2_ for 4 days until clear plaques formed. The plates were fixed in 4% formaldehyde solution for 2 hours and stained with 1% crystal violet.

### Focus-forming assay

The FFA was performed on C6/36 mosquito cells as described previously^21^. In brief, C6/36 cells were seeded into 12-well plates 12-16 h prior to the assay. A 25 μl volume of each sample was used for 10-fold serial dilutions. For each dilution, 200 μl were added to 12-well plates with 90% confluent cells. The cells were incubated at 30°C with 5% CO_2_ for 1 h with gentle rocking every 15 min. After a 5-day incubation, plates were fixed with methanol / acetone (1:1), washed by PBS, and blocked with 2% FBS/PBS before overnight incubation with mouse anti-ZIKV antibody (4G2). Plates were washed and incubated with goat anti-mouse secondary antibody conjugated to horseradish peroxidase (KPL, Gaithersburg, MD, USA), then washed and developed with aminoethyl carbazole solution (Enzo Diagnostics, Farmingdale, NY, USA) prepared according to the manufacturer’s protocol for detection of infection foci.

### Construction of ZIKV mutant infectious clones

The ZIKV FSS13025 full-length cDNA infectious clone pFLZIKV-FSS^41^ (referred as FSS-WT in text) was used as the backbone for engineering FSS-T106A, FSS-V1A, FSS-A188V, FSS-M872V and FSS-4M mutants. The ZIKV Dakar 41525 full-length cDNA infectious clone pFLZIKV-DK (referred as DK-WT in text) was used as the backbone for engineering DK-A106T, DK-A1V, DK-V188A, DK-V872M and DK-4M mutants. Standard overlapping PCR was performed to amplify the cDNA fragment between unique restriction enzyme sites that contained the corresponding mutations. Afterward, the fragments were cloned into pFLZIKV-FSS or pFLZIKV-DK plasmids and propagated in E. coli strain Top 10 (ThermoFisher Scientific, Framingham, MA, USA). All restriction enzymes were purchased from New England BioLabs (Massachusetts, USA). The plasmids were validated by restriction enzyme digestion and Sanger DNA sequencing. All primers were synthesized from Integrated DNA Technologies and are available upon request.

### RNA transcription, electroporation and virus recovery

The full-length cDNA clone plasmids of all ZIKVs were linearized with restriction enzyme ClaI prior to RNA transcription. The linearized DNAs were purified by phenol-chloroform extraction and ethanol precipitation. RNA transcripts of ZIKVs were synthesized using the T7 mMessage mMachine kit (Ambion, California, USA) *in vitro.* The quantity and quality of RNAs were verified by Spectrophotometry (DS-11, DeNovix, Delaware, USA) and Agarose gel electrophoresis. For RNA transfection, 10 μg of transcribed RNA were electroporated into 8×10^6^ Vero cells using the Gene Pulser Xcell™ Electroporation Systems (Bio-rad, California, USA) under as described previously^41^. The viral culture supernatant was harvested when obvious cytopathic effects (CPE) of the transfected Vero cells were observed, and the titers of all ZIKVs were measured on Vero cells using plaque assays^41^.

### Statistics

Animals were randomly allocated to different groups. Mosquitoes that died before measurement were excluded from the analysis. The investigators were not blinded to the allocation during the experiments or to the outcome assessment. No statistical methods were used to predetermine the sample size. Descriptive statistics are provided in the figure legends. Linear regression analysis was used to assess the correlation between WT/Mutant ratios by Sanger sequencing and WT/Mutant ratios by PFU. Analysis was performed in Prism version 7.03 (GraphPad, San Diego, 440 CA).

For virus competition experiments in mosquitoes, A129 mice and human primary cells, relative replicative fitness values for viruses in each sample were analyzed according to w=(f0/i0), where i0 is the initial ratio of one competitor and f0 is the final ratio after competition. Sanger sequencing (initial timepoint T0) counts for each virus strain being compared (for example, wildtype versus mutant strains) were based upon average counts over several replicate samples of inocula or bloodmeals per experiment (effectively averaging the replicates), and post-infection (timepoint T1) counts were taken from samples of individual subjects, such that for each strain the ratio between counts at T0 and T1 (T1/T0) reflects a measure of a sample from a single mosquito, mouse or cell culture well in a specific experiment for that strain. Typically, multiple experiments were performed, so that f0/i0 was clustered by experiment. To model f0/i0, the ratio T0/T1 was found separately for each subject in each strain group, log (base-10) transformed to an improved approximation of normality, and modeled by analysis of variance with relation to group, adjusting by experiment to control for clustering within experiment. Specifically, the model was of the form Log10_CountT1overCountT0 ~ Experiment + Group. Fitness ratios between the two groups [the model’s estimate of w=(f0/i0)] were assessed per the coefficient of the model’s Group term, which was transformed to the original scale as 10^coefficient. This modeling approach compensates for any correlation due to clustering within experiment similarly to that of corresponding mixed effect models, and is effective since the number of experiments was small. Statistical analyses were performed using R statistical software (R Core Team, 2019, version 3.6.1). In all statistical tests, two-sided alpha=.05. Catseye plots^42^, which illustrate the normal distribution of the model-adjusted means, were produced using the “catseyes” package^43^.

## Data availability

Extended Data and source data for generating main figures are available in the online version of the paper. Any other information is available upon request.

## Acknowledgments

S.C.W was supported by NIH grants AI120942 and 121452. P.-Y.S. was supported by NIH grants AI142759, AI127744, and AI136126, and awards from the Kleberg Foundation, John S. Dunn Foundation, Amon G. Carter Foundation, Gilson Longenbaugh Foundation, and Summerfield Robert Foundation. J. L. was supported by the McLaughlin Fellowship Fund. We thank Aaron Brault for helpful suggestions for the manuscript.

## Author Contributions

S.C.W., J.L., Y.L., C.S., and P.-Y.S. designed the experiments and/or wrote the manuscript; J.L. and Y.L. performed the majority of the experiments and analyzed the data; S.C., B.T.D.N., and P.-Y.S. developed the reverse genetic system for ZIKV strains; R.M. and N.V. provided the mosquito colonies; R.M. assisted in mosquito collection; S.L.H. assisted in designing the competition assay; G.H.R. and S.R.A provide the A129 mice colonies; C.R.A. assisted with statistics analyses; K.P. provided ZIKV parental strains and contributed experimental suggestions. All authors reviewed, critiqued and provided comments on the text.

## Extended Data Figures and Tables

**Extended Data Fig. 1.**
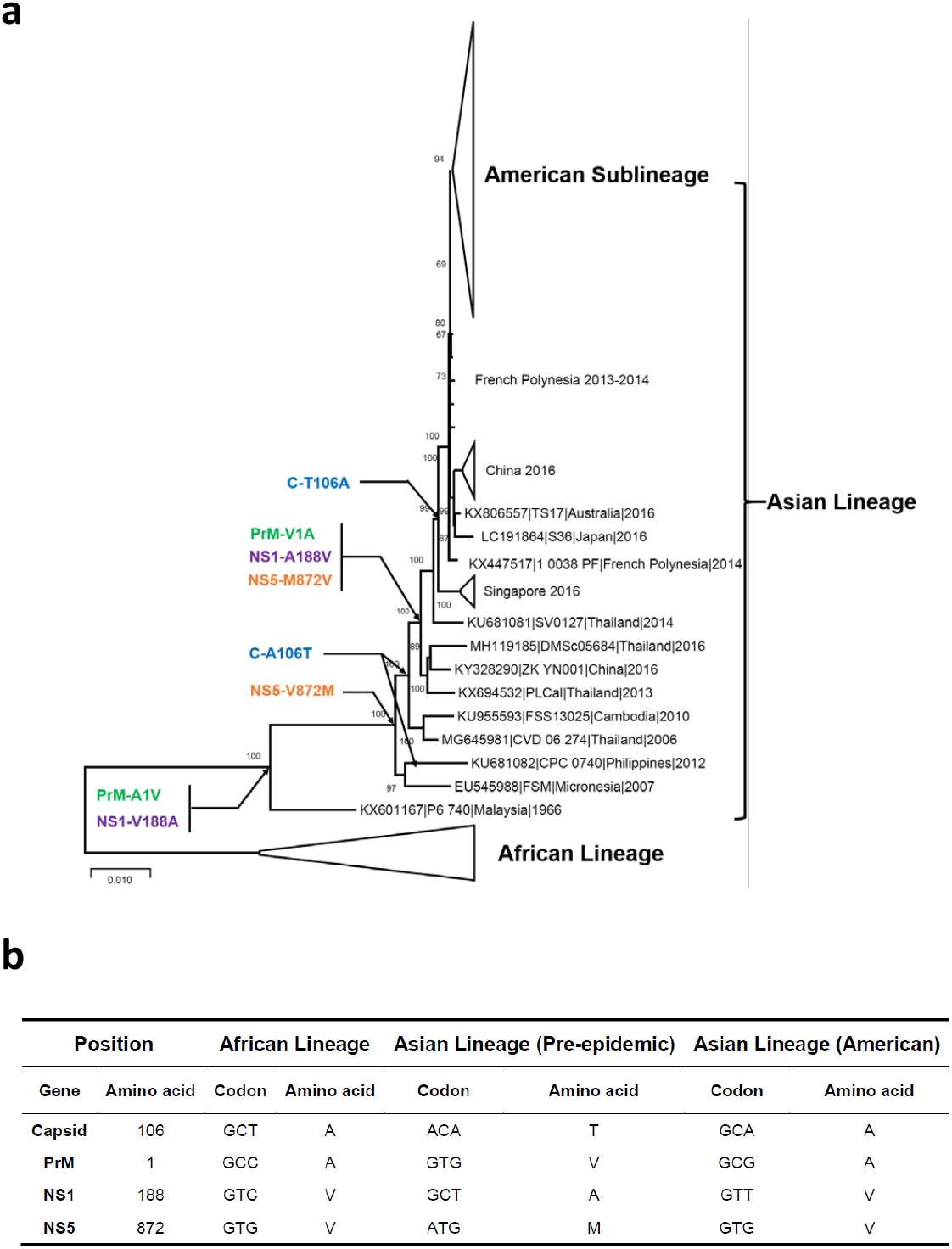
Phylogenetic analysis of representative ZIKV strains. **a**, Phylogenetic tree of representative Zika virus (ZIKV) strains based on 73 complete open reading frame sequences. The evolutionary distances were computed using the Maximum Composite Likelihood method. The percentage of replicate trees in which the associated taxa clustered together in the bootstrap tests were shown next to the branches (1000 replicates). Evolutionary analyses were performed in MEGA X, by using the Maximum Likelihood method. **b**, Four reversion directly reverting mutations compared among representative African, Asian and American lineage strains.

**Extended Data Fig. 2.**
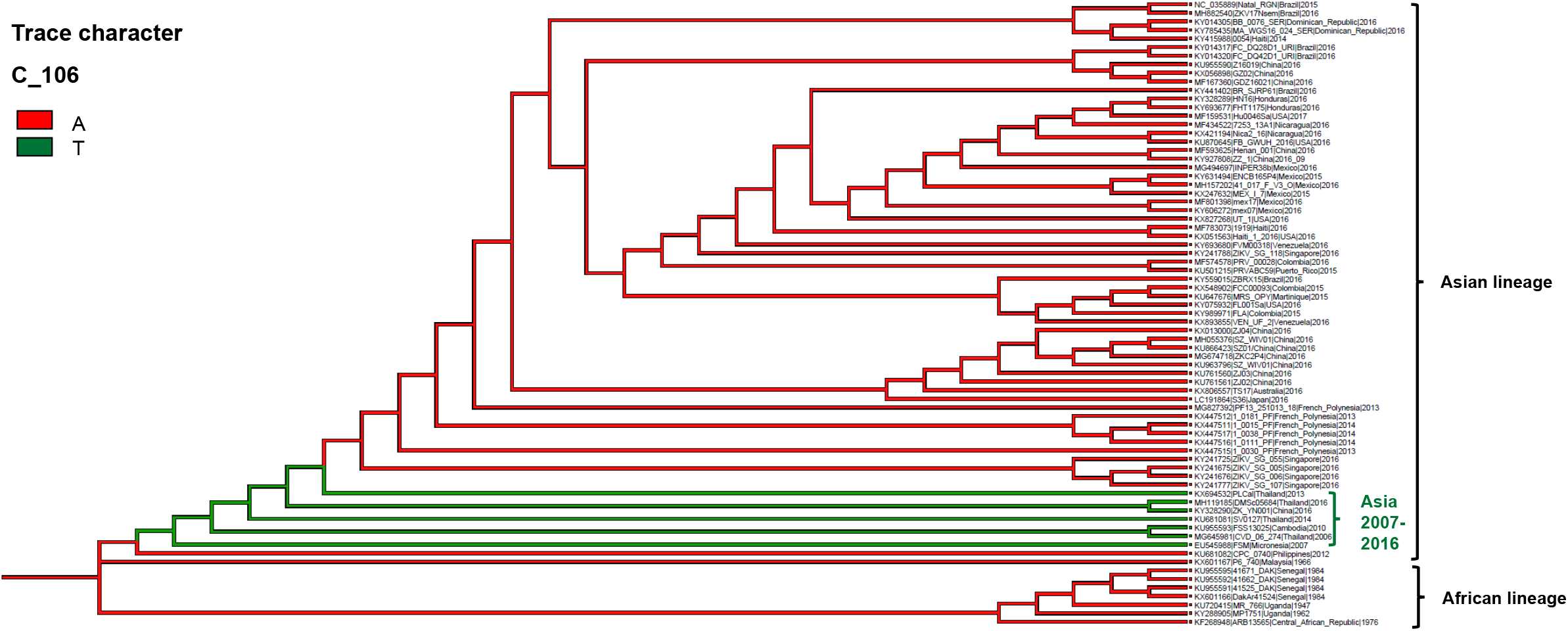
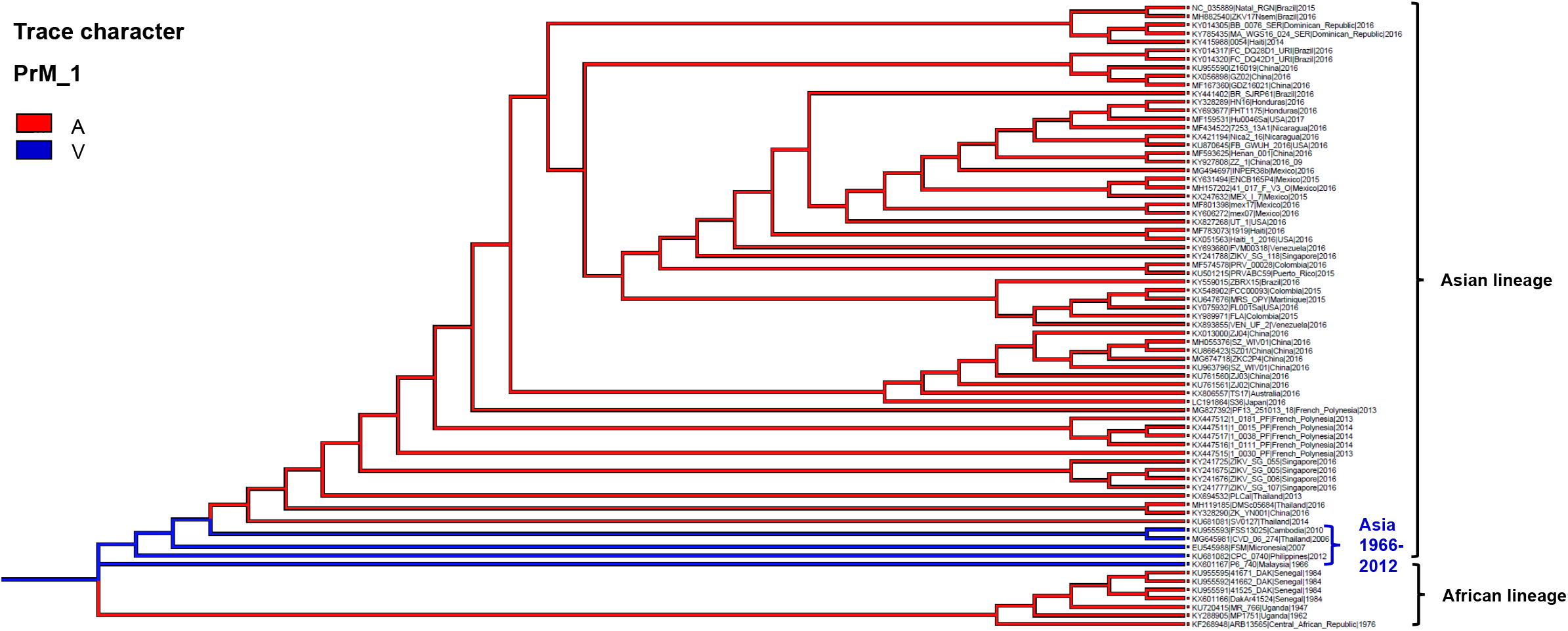
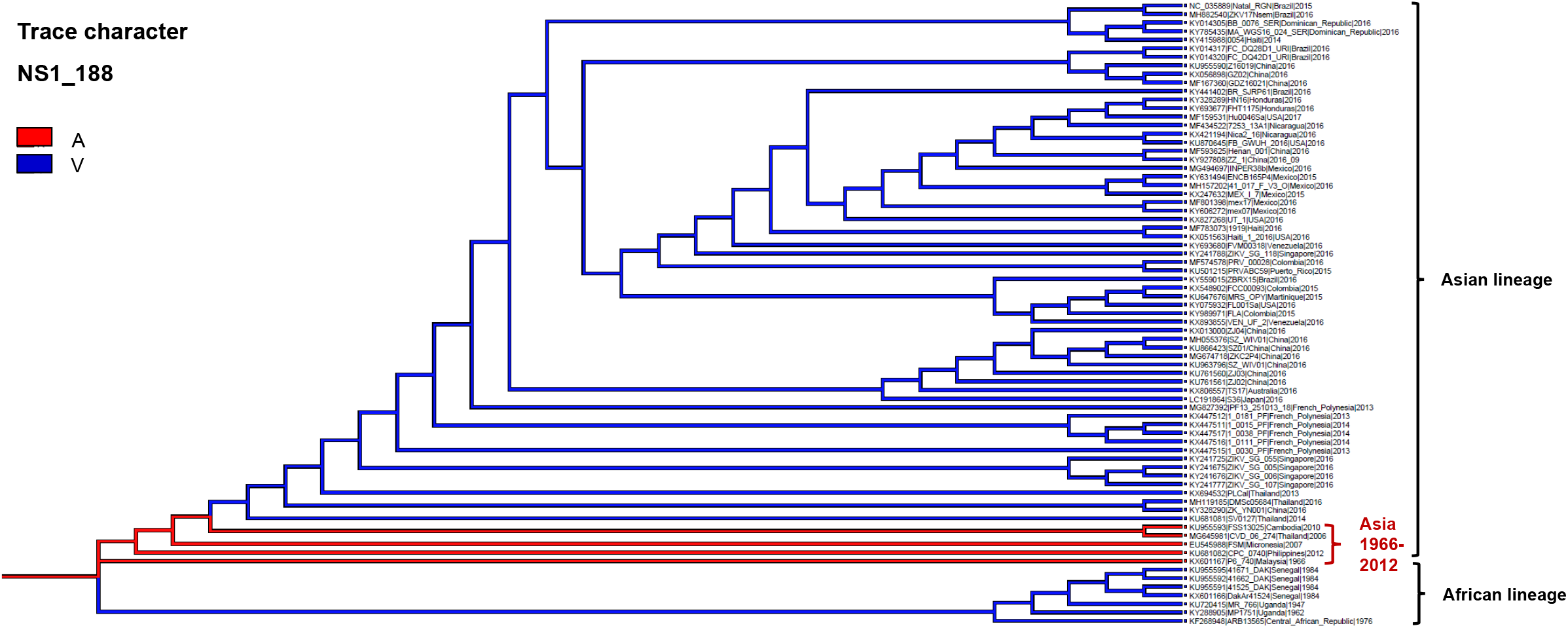
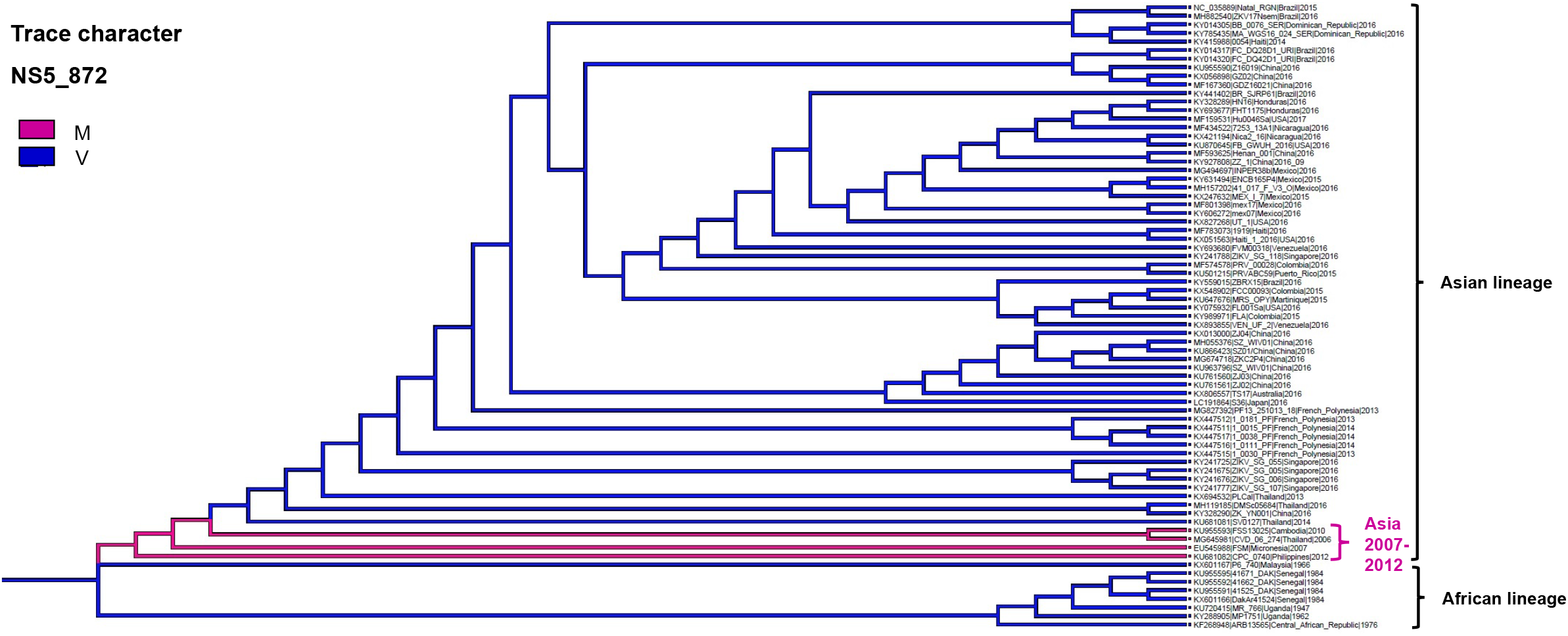
The trace history of four directly reverting amino acids in the evolutionary tree of Zika virus. Trace history of four ZIKV amino acids performed in Mesquite (Version 3.61, http://www.mesquiteproject.org). These include: **a**. Amino acid 106 substitution of the capsid protein; **b**. Amino acid 1 of the pre-membrane (prM) protein; **c**. amino acid 188 of the nonstructural protein 1 (NS1), and; **d**. amino acid 872 of the nonstructural protein 5 (NS5) based on 73 complete zika virus open reading frame sequences.

**Extended Data Fig. 3.**
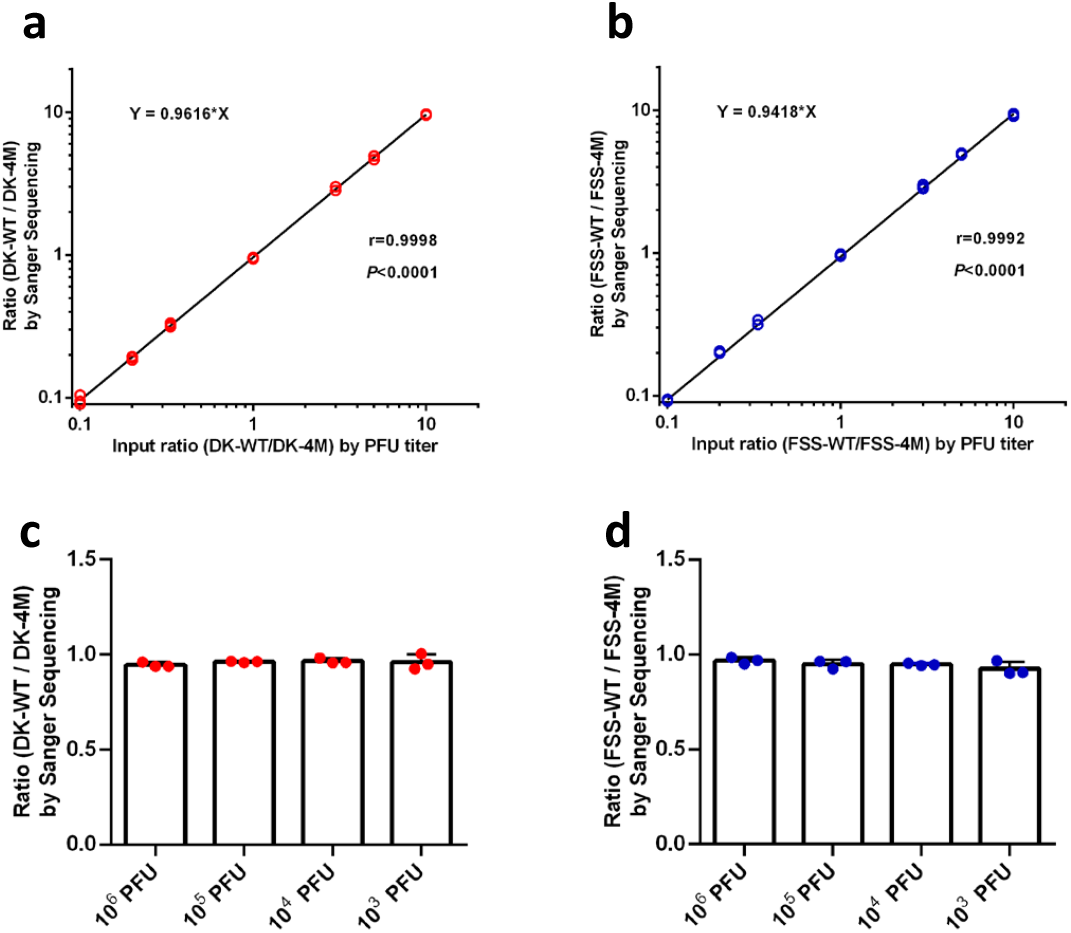
The consistency and accuracy validation of competition assay by Sanger sequencing. **a**, **b**, The correlation between PFU input ratios and output ratios determined by Sanger sequencing of RT-PCR amplicons. The DK-WT/DK-4M (**a**) or FSS-WT/FSS-4M(**b**) ZIKVs were mixed at different ratios of 10:1, 5:1, 3:1, 1:1, 1:3, 1:5, 1:10 based on the PFU titer. The total RNA of these mixed virus was isolated and transcribed by RT-PCR. The ratios of DK-WT/DK-4M and FSS-WT/FSS-4M were calculated based on the peak heights by Sanger sequencing. Data were analyzed by linear regression with correlation coefficients (r) and significance (p). **c**, **d**, The ratio of wt/mutant virus mixture calculated by Sanger sequencing was consistent when using virus mixture that ranges from high to low titer. The DK-WT/DK-4M (**c**) or FSS-WT/FSS-4M(**d**) ZIKVs were mixed at a PFU ratio of 1:1. The total titers of the mixed viruses were 10^6^, 10^5^, 10^4^ and 10^3^ PFU. The total RNA of these mixed virus was isolated and transcribed by RT-PCR. The ratios of DK-WT/DK-4M and FSS-WT/FSS-4M were calculated based on the peak heights by Sanger sequencing.

**Extended Data Fig. 4.**
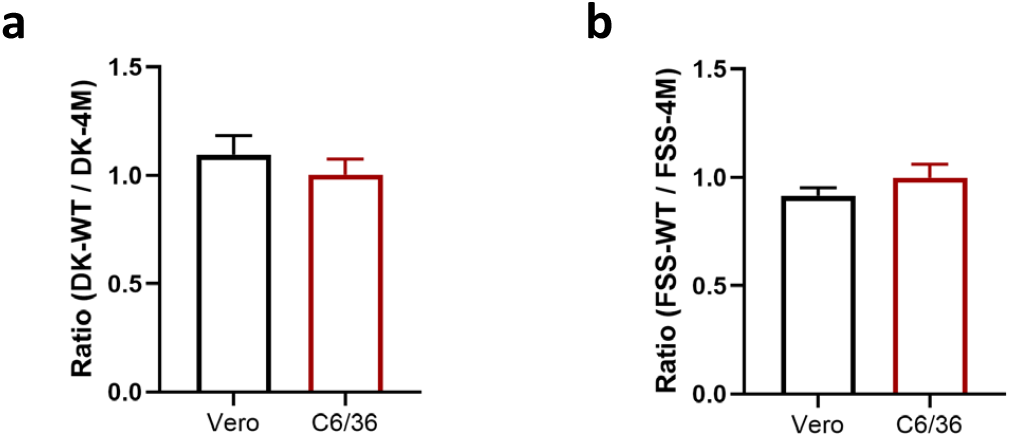
ZIKV WT and 4M mutant strain had a similar initial ratio when titrated on mammalian or insect cells. The initial ratio of DK-WT/DK-4M (**a**) and FSS-WT/FSS-4M (**b**) were calculated by the virus titer determined on Vero and C6/C36 cells. The titers of the WT and mutant viruses were determined by FFA (Focus-forming assay) on the cells before they were mixed.

**Extended Data Fig. 5.**
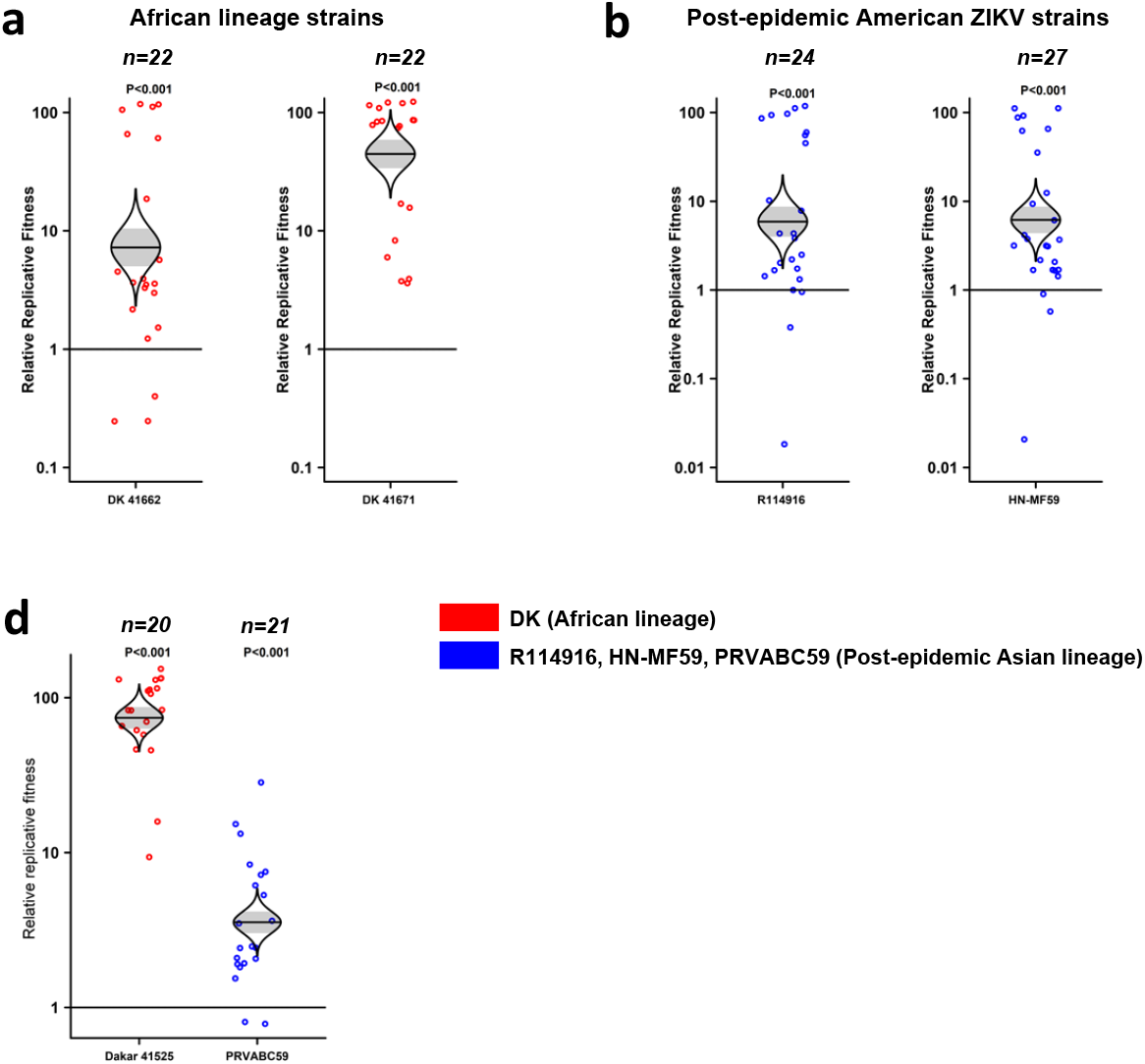
Fitness comparisons with additional Zika virus strains Dominican Republic *Aedes aegypti* mosquitoes. **a**, The fitness of 2 African ZIKV strains (DK 41662) and (DK 41671) in competition with the Asian FSS13025 strain in mosquitoes. **b**, The fitness of 2 American ZIKV strains (R114916) and (HN-MF59) in competition with the FSS13025 Asian strain in mosquitoes. **c.** The fitness of ZIKV African strain Dakar 41525, Asian pre-epidemic strain FSS13025 strain and American strain PRVABC59 in a low generation (F6) colonized *A. aegypti* strain from the Dominican Republic. Each point represents a single mosquito or mouse sample **a**-**c**, The distribution of the model-adjusted means is illustrated by catseye plots with shaded +/− standard error overlaid by scatterplots of subject measures; scatterplots have been randomly jittered horizontally for clarity, and are shown on the log (base-10) scale such that comparisons are against a null value of 1.

**Extended Data Fig. 6.**
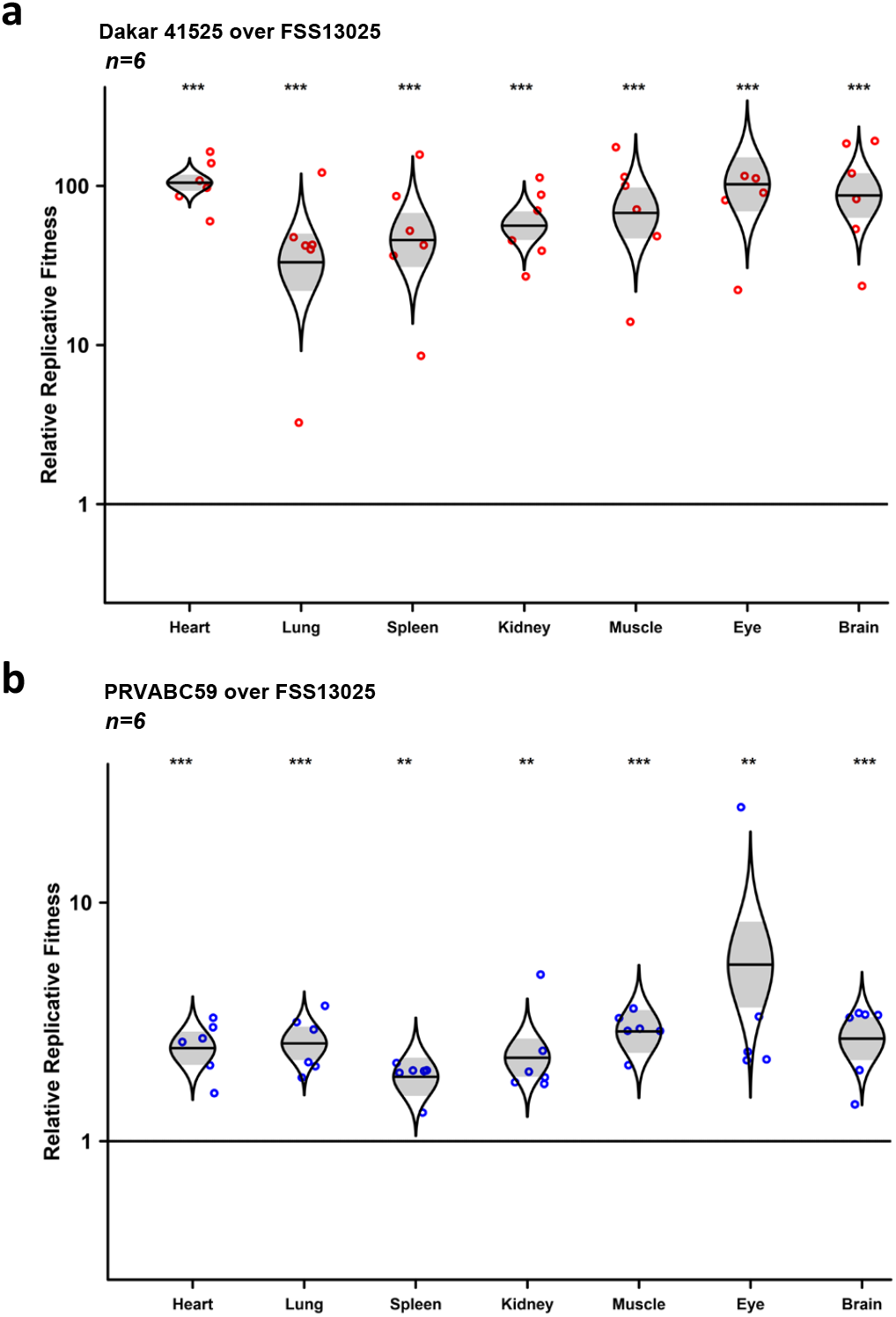
Fitness comparisons of African and post-epidemic against a pre-epidemic ZIKV strains in different organs of A129 mice. **a**, Fitness comparison between African (Dakar 41525) and Asian pre-epidemic (FSS13025) ZIKV strains in different organs of A129 mice after 8 days of infection, when viremia had ended. **b**, The fitness comparison between Asian pre-epidemic (FSS13025) and American (PRVABC59) ZIKV strains in different organs of A129 mice after 8 days of infection, when viremia had ended. Each point represents a single mouse sample **a**, **b**, The distribution of the model-adjusted means is illustrated by catseye plots with shaded +/− standard error overlaid by scatterplots of subject measures; scatterplots have been randomly jittered horizontally for clarity, and are shown on the log (base-10) scale such that comparisons are against a null value of 1. * *P* < 0.05, ** *P* < 0.01, *** *P* < 0.001.

**Extended Data Fig. 7.**
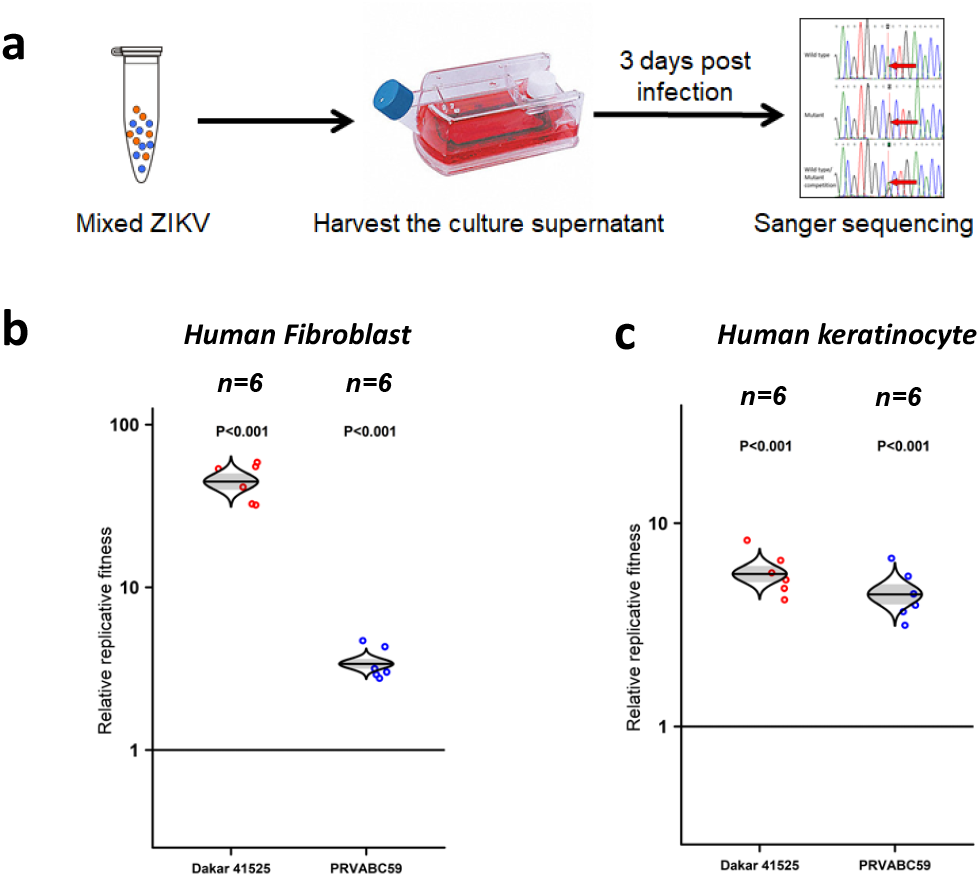
Fitness comparison of African and post-epidemic against a pre-epidemic ZIKV stain in human primary cells. **a**, Schematic representation of the study design. The mixed ZIKVs were inoculated into human fibroblast and keratinocyte cells. The RNAs in the culture supernatant were isolated, amplified by RT-PCR, Sanger-sequenced 3 days post infection. **b**, **c**, The fitness of ZIKV African strain (Dakar 41525), Asian pre-epidemic (FSS13025) strain and American strain (PRVABC59) in human primary fibroblast (**b**) and keratinocyte (**c**) cells. Each point represents a single culture sample The distribution of the model-adjusted means is illustrated by catseye plots with shaded +/− standard error overlaid by scatterplots of subject measures; scatterplots have been randomly jittered horizontally for clarity, and are shown on the log (base-10) scale such that comparisons are against a null value of 1.

**Extended Data Fig. 8.**
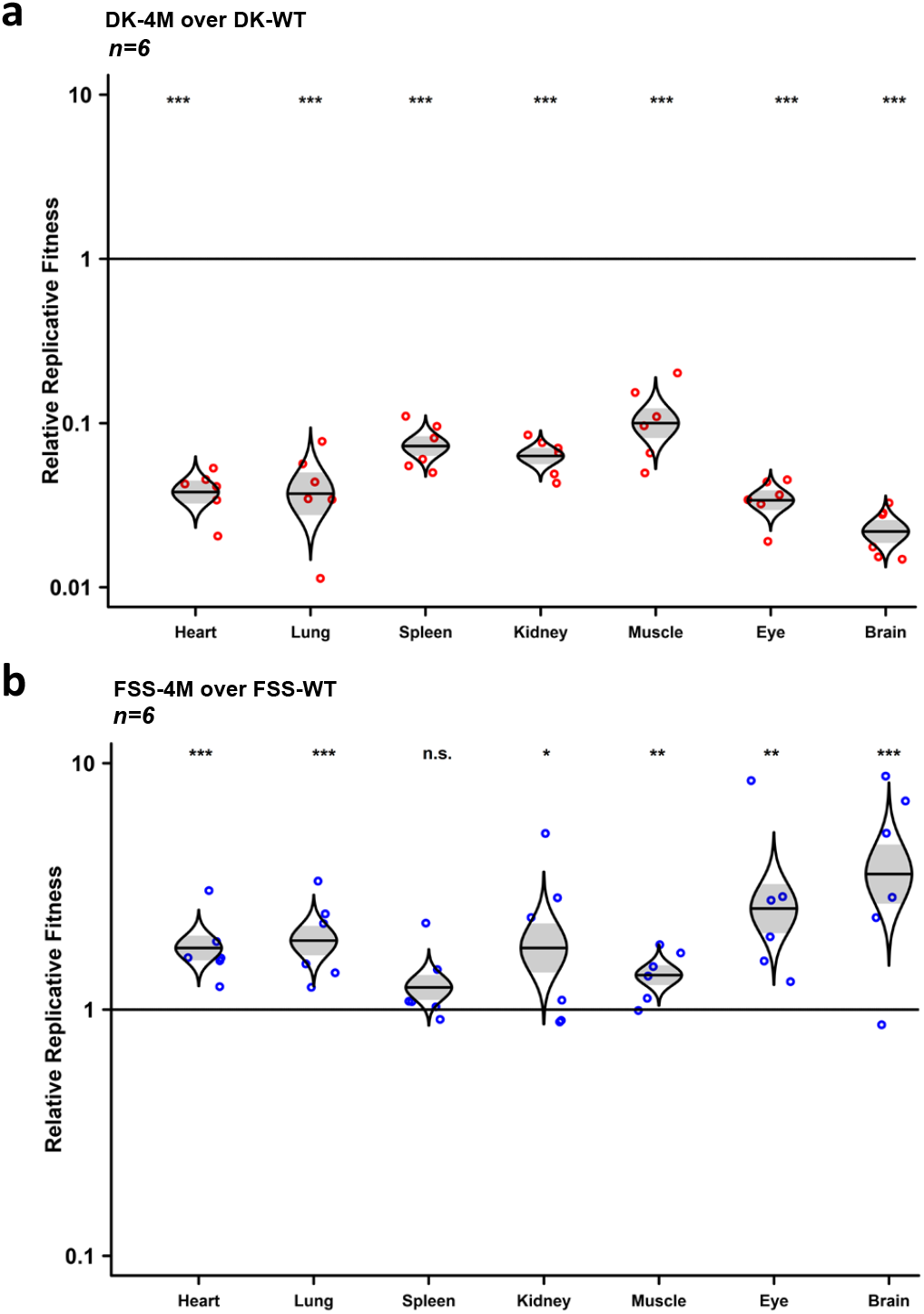
Fitness comparisons of Dakar-4 amino-acid mutant and FSS13025-4 amino-acid mutant against wild-type viruses in different organs of A129 mice. **a**, Fitness comparison between Dakar 4-amino acid mutant (DK-4M) and the wild-type strain in different organs of A129 mice. **b**, The fitness comparison between FSS 4 amino-acid mutant (FSS-4M) and wild-type strain in different organs of A129 mice. Each point represents a single mouse sample. **a**, **b**, The distribution of the model-adjusted means is illustrated by catseye plots with shaded +/− standard error overlaid by scatterplots of subject measures; scatterplots have been randomly jittered horizontally for clarity, and are shown on the log (base-10) scale such that comparisons are against a null value of 1. * *P* < 0.05, ** *P* < 0.01, *** *P* < 0.001, n.s. not significant.

**Extended Data Fig. 9.**
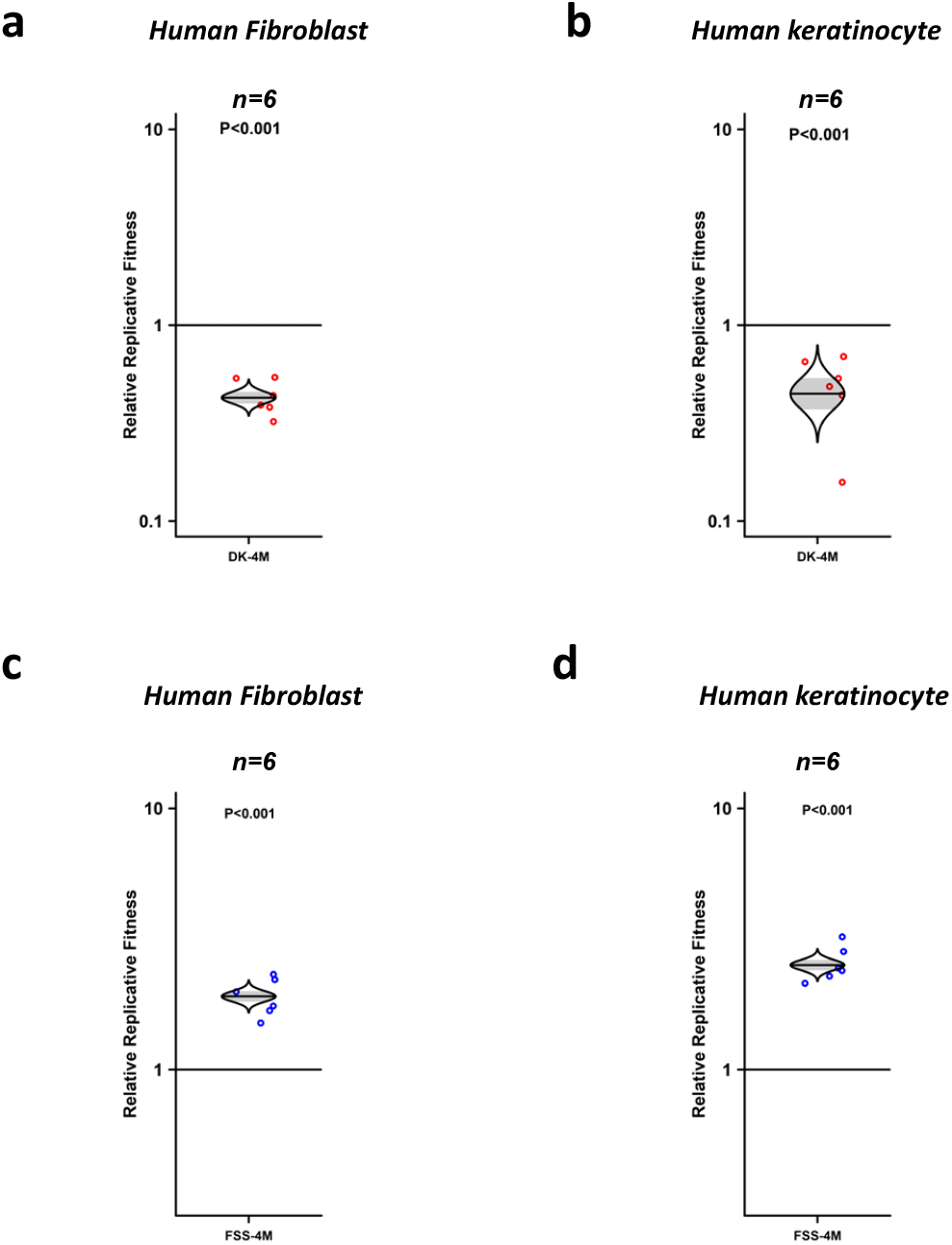
Fitness comparisons of Dakar-4 amino-acid mutant and FSS13025-4 amino-acid mutant versus wild-type ZIKV strains in human primary cells. **a**, **b**, Fitness comparison between Dakar 4-amino acid mutant (DK-4M) and wild-type strain in human primary fibroblast (**a**) and keratinocyte (**b**) cells. **c**, **d**, The fitness comparison between FSS13025 4-amino acid mutant (FSS-4M) and wild-type strain in human primary fibroblast (**c**)and keratinocyte (**d**) cells. Each point represents a single culture sample. **a**-**d**, The distribution of the model-adjusted means is illustrated by catseye plots with shaded +/− standard error overlaid by scatterplots of subject measures; scatterplots have been randomly jittered horizontally for clarity, and are shown on the log (base-10) scale such that comparisons are against a null value of 1.

**Extended Data Fig. 10.**
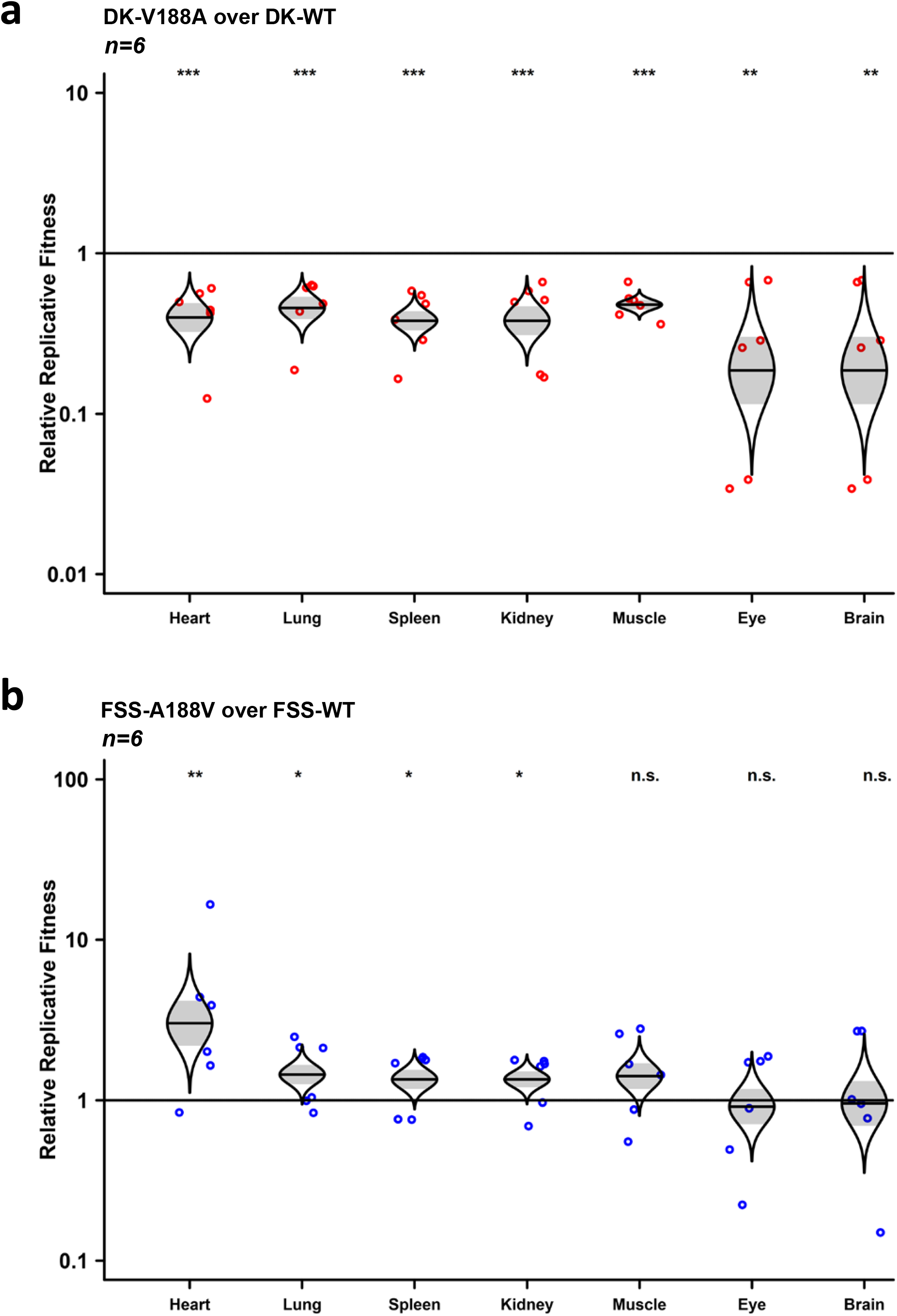
Fitness comparison between NS1-188 mutants against wild-type ZIKV strains in different organs of A129 mice. **a**, Fitness comparison between Dakar NS1-V188A mutant and wild-type strain in different organs of A129 mice. **b**, Fitness comparison between FSS13025 NS1-A188V mutant and wild-type strain in different organs of A129 mice. Each point represents a single mouse sample **a**, **b**, The distribution of the model-adjusted means is illustrated by catseye plots with shaded +/− standard error overlaid by scatterplots of subject measures; scatterplots have been randomly jittered horizontally for clarity, and are shown on the log (base-10) scale such that comparisons are against a null value of 1. * *P* < 0.05, ** *P* < 0.01, *** *P* < 0.001, n.s. not significant.

**Extended Data Fig. 11.**
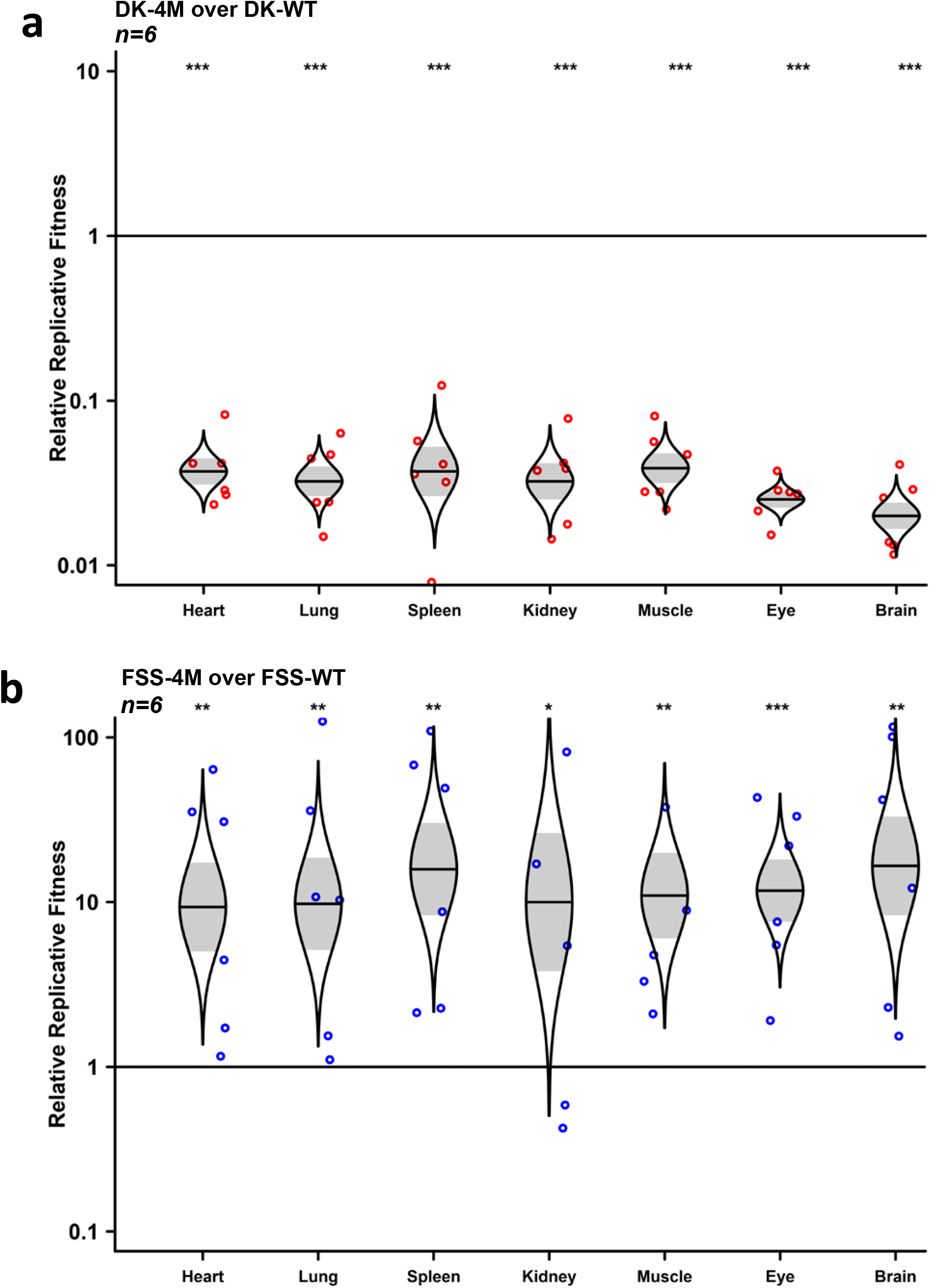
Fitness comparison between African-4 amino-acid mutant and FSS13025-4 amino-acid mutant against wild-type viruses during mosquito-mouse-mosquito transmission cycle. **a**, Fitness comparison between Dakar 4-amino acid mutant (DK-4M) and wild-type strain in different organs of A129 mice after being bitten by infected mosquitoes. **b**, Fitness comparison between FSS 4-amino acid mutant (FSS-4M) and wild-type strain in different organs of A129 mice biting by infected mosquitoes. Each point represents a single mosquito or mouse sample **a**, **b**, The distribution of the model-adjusted means is illustrated by catseye plots with shaded +/− standard error overlaid by scatterplots of subject measures; scatterplots have been randomly jittered horizontally for clarity, and are shown on the log (base-10) scale such that comparisons are against a null value of 1. * *P* < 0.05, ** *P* < 0.01, *** *P* < 0.001.

**Extended Data Table 1.**
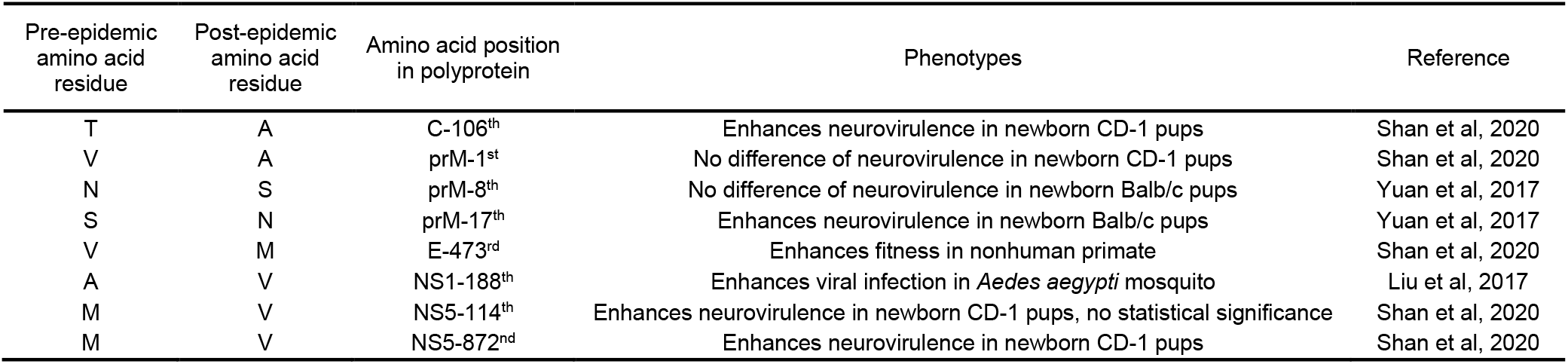
Summary of amino acid differences between recent, pre-epidemic Asian and post-epidemic Asian/American lineage ZIKVs. Phenotypes assigned to these amino acid substitutions are summarized.

| Pre-epidemic<br>amino acid<br>residue | Post-epidemic<br>amino acid<br>residue | Amino acid position<br>in polyprotein | Phenotypes | Reference |
| --- | --- | --- | --- | --- |
| T | A | C-106<sup>th</sup> | Enhances neurovirulence in newborn CD-1 pups | Shan et al, 2020 |
| V | A | prM-1<sup>st</sup> | No difference of neurovirulence in newborn CD-1 pups | Shan et al, 2020 |
| N | S | prM-8<sup>th</sup> | No difference of neurovirulence in newborn Balb/c pups | Yuan et al, 2017 |
| S | N | prM-17<sup>th</sup> | Enhances neurovirulence in newborn Balb/c pups | Yuan et al, 2017 |
| V | M | E-473<sup>rd</sup> | Enhances fitness in nonhuman primate | Shan et al, 2020 |
| A | V | NS1-188<sup>th</sup> | Enhances viral infection in *Aedes aegypti* mosquito | Liu et al, 2017 |
| M | V | NS5-114<sup>th</sup> | Enhances neurovirulence in newborn CD-1 pups, no statistical significance | Shan et al, 2020 |
| M | V | NS5-872<sup>nd</sup> | Enhances neurovirulence in newborn CD-1 pups | Shan et al, 2020 |

**Extended Data Table 2.**
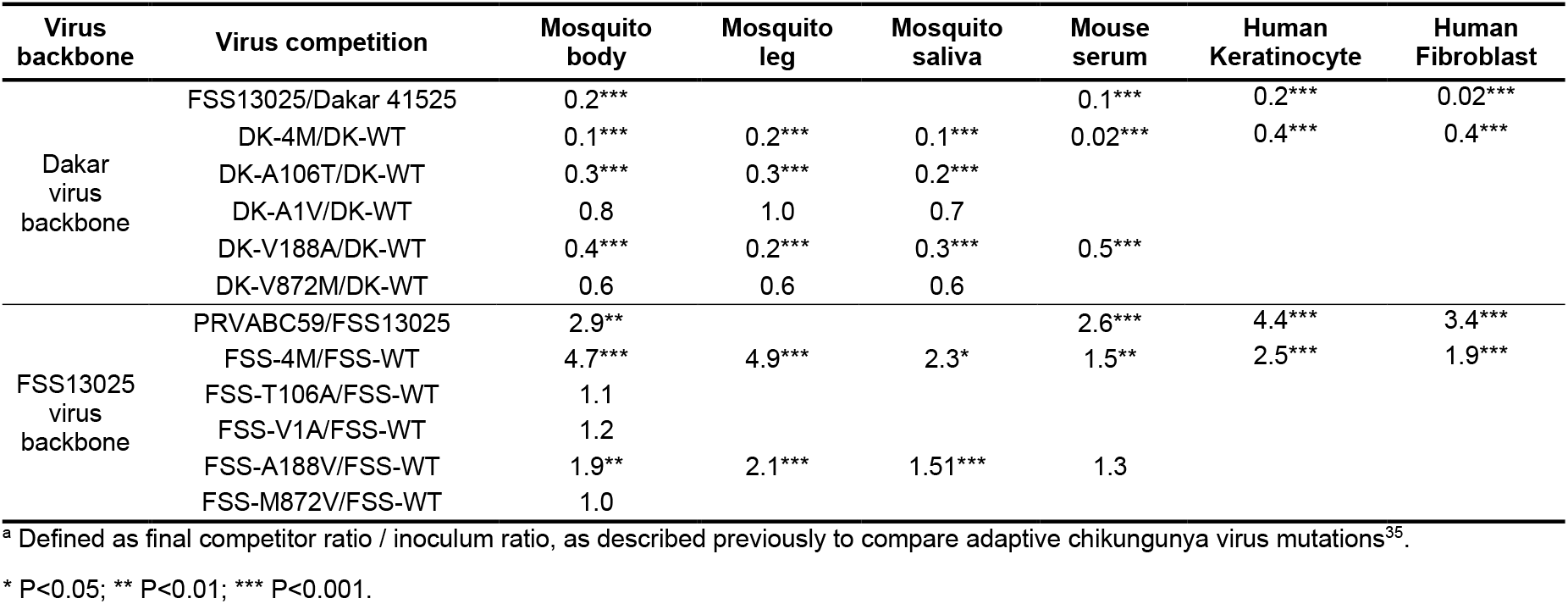
The relative replicative fitness of ZIKV mutant strains in mosquitoes, mice and human primary cells. The relative replicative fitness of ZIKV mutant strains in mosquitoes, mice and human primary cells, derived from the data in the figures, are presented as the final ratio divided by the initial ratio of the two competing viruses, as used previously to compare adaptive chikungunya virus mutations^38^. P values are based on differences from equal fitness or a relative fitness value of 1.

| Virus backbone | Virus competition | Mosquito body | Mosquito leg | Mosquito saliva | Mouse serum | Human Keratinocyte | Human Fibroblast |
| --- | --- | --- | --- | --- | --- | --- | --- |
| Dakar virus backbone | FSS13025/Dakar 41525 | 0.2*** |  |  | 0.1*** | 0.2*** | 0.02*** |
|  | DK-4M/DK-WT | 0.1*** | 0.2*** | 0.1*** | 0.02*** | 0.4*** | 0.4*** |
|  | DK-A106T/DK-WT | 0.3*** | 0.3*** | 0.2*** |  |  |  |
|  | DK-A1V/DK-WT | 0.8 | 1.0 | 0.7 |  |  |  |
|  | DK-V188A/DK-WT | 0.4*** | 0.2*** | 0.3*** | 0.5*** |  |  |
|  | DK-V872M/DK-WT | 0.6 | 0.6 | 0.6 |  |  |  |
| FSS13025 virus backbone | PRVABC59/FSS13025 | 2.9** |  |  | 2.6*** | 4.4*** | 3.4*** |
|  | FSS-4M/FSS-WT | 4.7*** | 4.9*** | 2.3* | 1.5** | 2.5*** | 1.9*** |
|  | FSS-T106A/FSS-WT | 1.1 |  |  |  |  |  |
|  | FSS-V1A/FSS-WT | 1.2 |  |  |  |  |  |
|  | FSS-A188V/FSS-WT | 1.9** | 2.1*** | 1.51*** | 1.3 |  |  |
|  | FSS-M872V/FSS-WT | 1.0 |  |  |  |  |  |

<sup>a</sup> Defined as final competitor ratio / inoculum ratio, as described previously to compare adaptive chikungunya virus mutations<sup>35</sup>.
\* P<0.05; \*\* P<0.01; \*\*\* P<0.001.

**Extended Data Table 3.**
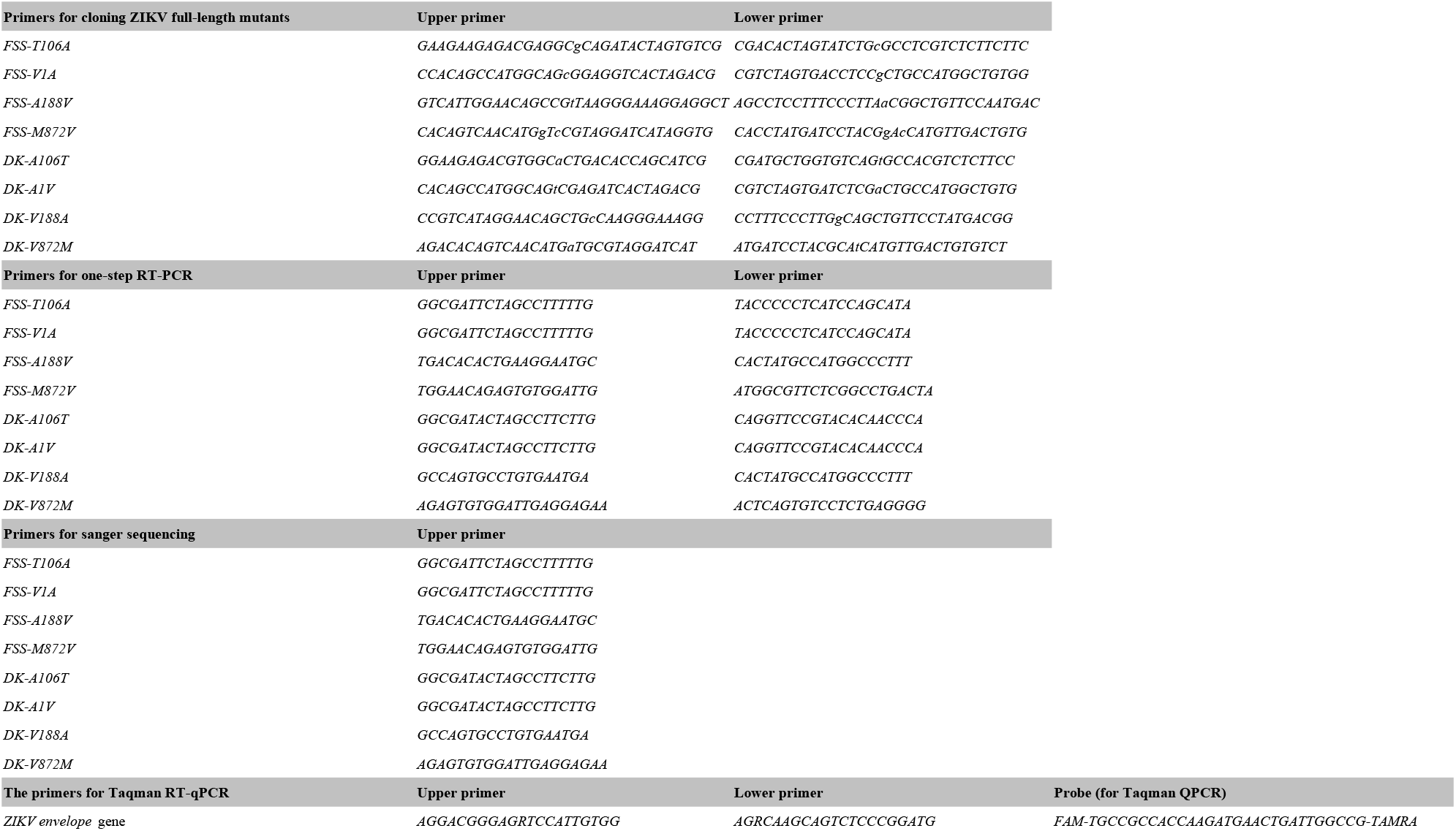
Primers and probes for gene cloning, qPCR, RT-PCR and Sanger sequencing.

